# HSF2BP Negatively Regulates Homologous Recombination in DNA Interstrand Crosslink Repair in Human Cells by Direct Interaction With BRCA2

**DOI:** 10.1101/438945

**Authors:** Inger Brandsma, Koichi Sato, Sari E. van Rossum-Fikkert, Marcel Reuter, Hanny Odijk, Nicole Verkaik, Nathalie van den Tempel, Anneke B. Oostra, Dick H. W. Dekkers, Karel Bezstarosti, Jeroen A. A. Demmers, Joyce Lebbink, Claire Wyman, Josephine C. Dorsman, Dik C. van Gent, Puck Knipscheer, Roland Kanaar, Alex N. Zelensky

## Abstract

The tumor suppressor BRCA2 is essential for homologous recombination, replication fork stability and DNA interstrand crosslink (ICL) repair in vertebrates. We show that a functionally uncharacterized protein, HSF2BP, is involved in a novel, direct and highly evolutionarily conserved interaction with BRCA2. Although *HSF2BP* was previously described as testis-specific, we find it is expressed in mouse ES cells, in human cancer cell lines, and in tumor samples. Elevated levels of HSF2BP sensitize human cells to ICL-inducing agents (mitomycin C and cisplatin) and PARP inhibitors, resulting in a phenotype characteristic of cells from Fanconi anemia (FA) patients. We biochemically recapitulate the suppression of ICL repair and establish that excess HSF2BP specifically compromises homologous recombination by preventing BRCA2 and RAD51 loading at the ICL. As increased ectopic expression of HSF2BP occurs naturally, we suggest that it can be considered as a causative agent in FA and a source of cancer-promoting genomic instability.

## Introduction

Breast cancer associated protein 2 (BRCA2) has been the subject of intense research because of its role as a tumor suppressor and the mediator of homologous recombination (HR) (Prakash et al., 2015; Roy et al., 2012; Zelensky et al., 2014). Yet important questions concerning the roles of specific domains in BRCA2, such as its most conserved region (the DNA-binding domain) or internally disordered regions, relative importance of distinct activities of the protein, etc. remain open. A major breakthrough in understanding the molecular function of BRCA2 came from the identification of RAD51 as an interaction partner (Sharan et al., 1997). Interaction of BRCA2 with other proteins, known (Martinez et al., 2015) and yet undiscovered, will help address some of the open questions.

During DNA interstrand crosslink (ICL) repair several substrates for HR arise, beginning with maintenance and rearrangement of the replication fork, and up to accurate repair of the DNA double-strand break (DSB) that is created after nucleolytic disengagement of the crosslinked strands. Recognition of the lesion, damage response signaling and coordination of the multiple distinct enzymatic reactions that need to happen to restore genetic information, is performed by a large and still growing group of proteins from the Fanconi anemia (FA) pathway, which as of now comprises of 22 proteins, designated FANC*A*–FANC*W* (Auerbach, 2009; Inano et al., 2017; Knies et al., 2017; Kottemann and Smogorzewska, 2013).

The functional connection between HR and FA has been demonstrated in numerous studies, however the macromolecular interactions underlying it are less clear. Several studies postulated HR-FA protein-protein interactions exist, such as FANCD2-BRCA1 (Garcia-Higuera et al., 2001), FANCA-BRCA1 (Folias et al., 2002), FANCD2-BRCA2 (Hussain et al., 2004; Wang et al., 2004; Zhi et al., 2009), FANCG-BRCA2-FANCD2-XRCC3 (Hussain et al., 2003; Wilson et al., 2008; 2010). However, these are suggestions mostly based on co-immunoprecipitation and co-localization to DNA damage-induced focal accumulations of proteins, with evidence for direct binding limited to yeast two-hybrid analyses. Given the complexity and the properties of the proteins involved, indirect interaction via DNA, chromatin or intermediary proteins may explain the reported co-occurrence of FA and HR proteins in foci and precipitates. On the other hand, the direct interactions that do take place and are responsible for the coordination between early FA factors and HR may be dynamic, low affinity, dependent on post-translational modifications, and thus evade discovery. The FANCD2-FANCI complex was recently demonstrated to directly bind to RAD51 using purified proteins (Sato et al., 2016). Characterization of other such interactions would help to understand the engagement between upstream FA and downstream HR machineries and interpret the existing data.

In this report we describe a novel, direct and highly evolutionarily conserved interaction of BRCA2 with HSF2BP, a protein previously only identified in testis and not characterized functionally. We show that the interaction between BRCA2 and HSF2BP ectopically produced in human cancer cells specifically disrupts HR in the context of ICL repair and sensitizes cells to poly (ADP-ribose) polymerase inhibitors (PARPi), but does not affect all BRCA2-dependent HR reactions, thus creating a setting in which BRCA2 functions are separated.

## Results

### HSF2BP is a member of the BRCA complex in mES cells

We previously described efficient immunoprecipitation of BRCA2-GFP and most of the known BRCA2-interacting proteins from *Brca2^GFP/GFP^* knock-in mouse embryonic stem (mES) cells (Reuter et al., 2014), and the phenomenon of BRCA2 degradation induced by mild hyperthermia (Krawczyk et al., 2011). With these at hand we performed quantitative SILAC-based mass spectrometry experiments and identified HSF2BP as one of the proteins whose abundance in the BRCA2-GFP immunoprecipitate co-varies with that of BRCA2 upon hyperthermia treatment **[Fig 1A]**. HSF2BP was previously described as a testis-specific, *<u>h</u>*eat *<u>s</u>*hock *f*actor <u>2</u> (HSF2) *<u>b</u>*inding *<u>p</u>*rotein (Yoshima et al., 1998), and its association with BRCA2 or expression in tissues other than testis has not yet been reported. A reciprocal mass spectrometry experiment – GFP immunoprecipitation from *Hsf2bp^GFP/+^* knock-in mES cells that we engineered – confirmed the interaction **[Fig 1B]**. As a control in these experiments we used immunoprecipitation from *Rad51ap1^GFP/GFP^* knock-in cells, to ensure the interactions were not due to non-specific binding to a nuclear GFP-tagged low abundance DNA repair proteins (Modesti et al., 2007). The efficiency of BRCA2 immunoprecipitation via HSF2BP-GFP was remarkably high, similar to direct BRCA2-GFP immunoprecipitation, which is noteworthy as in our experience immunoprecipitation of large low abundance HR proteins is not straightforward. Furthermore, HSF2BP co-precipitated with PALB2, the partner and localizer of BRCA2 (Xia et al., 2006) from *Palb2^GFP/GFP^* knock-in mES cells **[Fig 1C]**.

**Figure 1.**
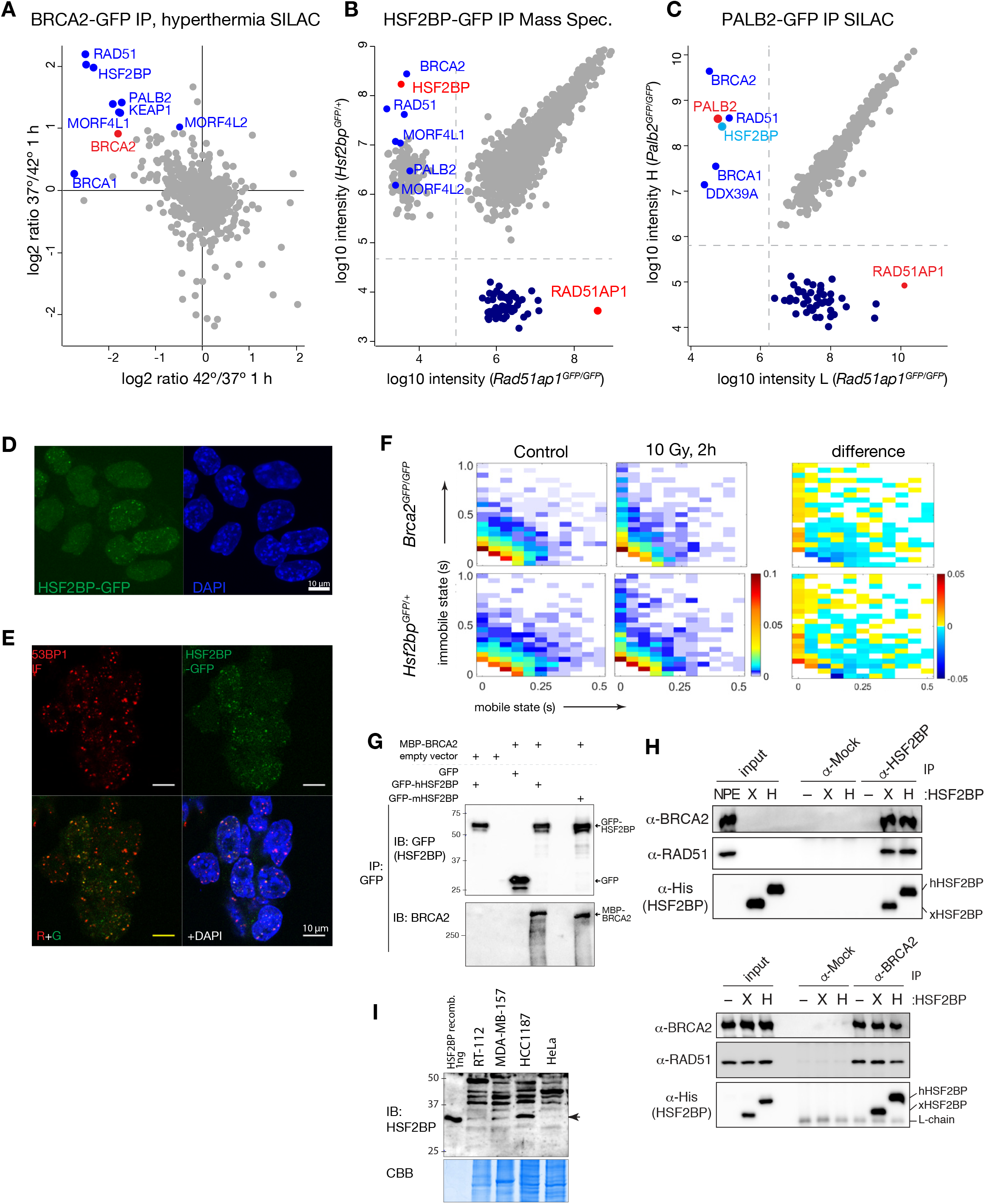
HSF2BP is a member of the BRCA complex. Identification of HSF2BP as BRCA2-interacting protein (**A**) Mass spectrometry analysis of anti-GFP immunoprecipitated proteins from Brca2GFP/GFP mES labelled by SILAC. Log2-transformed SILAC ratios from two separate (label-swap) experiments are plotted. One of the SILAC states in each experiment corresponds to 1 h incubation at 42 °C, which leads to BRCA2 degradation, the other to 37 °C control. Known members of BRCA complex are highlighted and labelled. (**B**) Proteins co-immunoprecipitated in two separate experiments with HSF2BP-GFP (vertical axis) or RAD51AP1-GFP (horizontal axis) were identified by mass-spectrometry. Combined peptide intensities (Log10-transformed) are plotted. Missing intensity values representing proteins identified only in one of the anti-GFP immunoprecipitates were replaced with values imputed from normal distribution with a downshift of 3 for visualization purpose only; their ranges are demarcated with dotted lines. (**C**) Proteins co-immunoprecipitating with PALB2-GFP and RAD51AP1-GFP from mES cells labelled by SILAC and quantified by mass-spectrometry; log10 transformed intensities are plotted, missing values imputed as in (B). (**D**) Hsf2bpGFP/+ cells were irradiated with 8 Gy, fixed, mounted in DAPI-containing media, and imaged by confocal microscopy. Maximum projection of direct GFP and DAPI fluorescence from a z-stack is shown. (**E**) Hsf2bpGFP/+ cells were irradiated with 8 Gy, fixed, processed for immunofluorescent staining with anti-53BP1 antibody (red channel) and mounted with DAPI. HSF2BP-GFP fluorescence was registered directly. (**F**) Mobility of single HSF2BP-GFP and BRCA2-GFP particles in live mES cells imaged by oblique illumination microscopy and analyzed using a single particle tracking routine as described (Reuter et al., 2014). Duration of mobile and immobile states under control conditions (left column) or 2 h after irradiation with 10 Gy (middle), and the difference between the two (right) are shown. (**G**) Interaction between hBRCA2 and HSF2BP revealed by anti-GFP immunoprecipitation from HEK293T cells co-expressing GPF-HSF2BP (human or mouse) and MBP-BRCA2. (**H**) Interaction between endogenous X. laevis BRCA2 and purified recombinant human (H) or X. laevis (X) HSF2BP revealed by co-immunoprecipitation from Xenopus egg extracts with or without recombinant HSF2BP added. (**I**) Immunoblot of HSF2BP in human cancer cell lines that were identified as having elevated HSF2BP mRNA levels. Whole cell lysates were separated by SDS-PAGE and analyzed by immunoblotting with anti-HSF2BP antibody SY8126.

HSF2BP-GFP in *Hsf2bp^GFP/+^* cells was predominantly nuclear and localized to ionizing radiation-induced foci readily detectable without pre-extraction **[Fig 1D]**. The foci co-localized with DSB marker 53BP1, but were observed in a fraction of cells **[Fig 1E]**, as expected given that HR activity, and specifically BRCA2 expression in mES cells (Reuter et al., 2014), is confined to S and G2 phases of cell cycle. We previously found using oblique illumination microscopy and single particle tracking that endogenous GFP-tagged BRCA2 diffuses in oligomeric clusters that become immobilized upon induction of DNA damage (Reuter et al., 2014). The same technique was applied to monitor and quantify HSF2BP-GFP diffusion in the nucleus, which revealed heterogeneous behavior very similar to that of BRCA2-GFP, including ionizing radiation-induced immobilization **[Fig 1F]**. Taken together, the immunoprecipitation mass spectrometry, and microscopy data suggest that HSF2BP and BRCA2 exist in a physiological complex in mES cells, which may be constitutive.

The interaction between HSF2BP and BRCA2 appears to be conserved, as in addition to the interaction between the mouse proteins, we detected association between their human orthologues **[Fig1 G]**. Furthermore, endogenous *X. laevis* BRCA2 interacted efficiently with both recombinant *X. laevis* HSF2BP (xlHSF2BP) and recombinant human HSF2BP **[Fig1 H]**. Most experiments described below were performed with the human proteins, and where mouse (mHSF2BP) and human (hHSF2BP) proteins were tested alongside in human cells, the observed effects were similar (see below). Thus HSF2BP-BRCA2 interaction is evolutionarily conserved, which is remarkable given the evolutionary distance between the species and indicates that the interaction is under strong selective pressure and thus functionally important.

We initially identified mHSF2BP in mES cells, could readily detect *HSF2BP* transcripts by RT-PCR in all human cancer cell lines we tested **[Fig S1A-D]**, and cloned the full-length hHSF2BP coding sequence (CDS) to produce the various expression constructs used in the study by RT-PCR from RNA isolated from human U2OS cells, which are from osteosarcoma rather than testis origin. By inspection of public RNA expression datasets (Harding et al., 2011; Petryszak et al., 2016; Shin et al., 2011; Tang et al., 2017), we found indications of high level expression of HSF2BP – in addition to testis – in mouse embryonic ovaries and some human cancer samples derived from brain, breast, ovarian and head and neck tumors **[Fig S1E]**. We tested three of the human cancer cell lines that had elevated *HSF2BP* expression levels (MDA-MB-157 (breast), HCC1187 (breast) and RT-112(bladder)), and could detect HSF2BP in all of them **[Fig 1I]**. These findings indicate that *HSF2BP* promoter can be activated in cancer cells of non-testicular origin. In conjunction with the BRCA2 interaction, this prompted us to explore its possible effects on homologous recombination.

### HSF2BP overproduction disrupts the FA pathway

Overexpression in human cancer samples and a serendipitous finding from initial RNAi experiments led us to question whether overproduction of HSF2BP may disrupt BRCA2 function in genome maintenance **[Fig 2]**. In the RNAi experiment we aimed to test the sensitivity of HSF2BP-depleted cells to genotoxic treatments and used GFP-HSF2BP as a surrogate measure of the knock-down efficiency **[Fig S2A]**. Unexpectedly, cells stably overproducing GFP-HSF2BP were hypersensitive to the DNA cross-linking agent mitomycin C (MMC), and the effect was reverted by RNAi **[Fig S2B]**. We further established that stable overproduction of untagged HSF2BP in human cells sensitizes them to MMC, cisplatin and PARP inhibitor olaparib **[Fig 2A-C, S2C]**. Plating efficiency and ionizing radiation sensitivity were not affected **[Fig 2D, S2D,E]**, indicating that sensitization was not due to reduced viability or major HR dysfunction.

**Figure 2.**
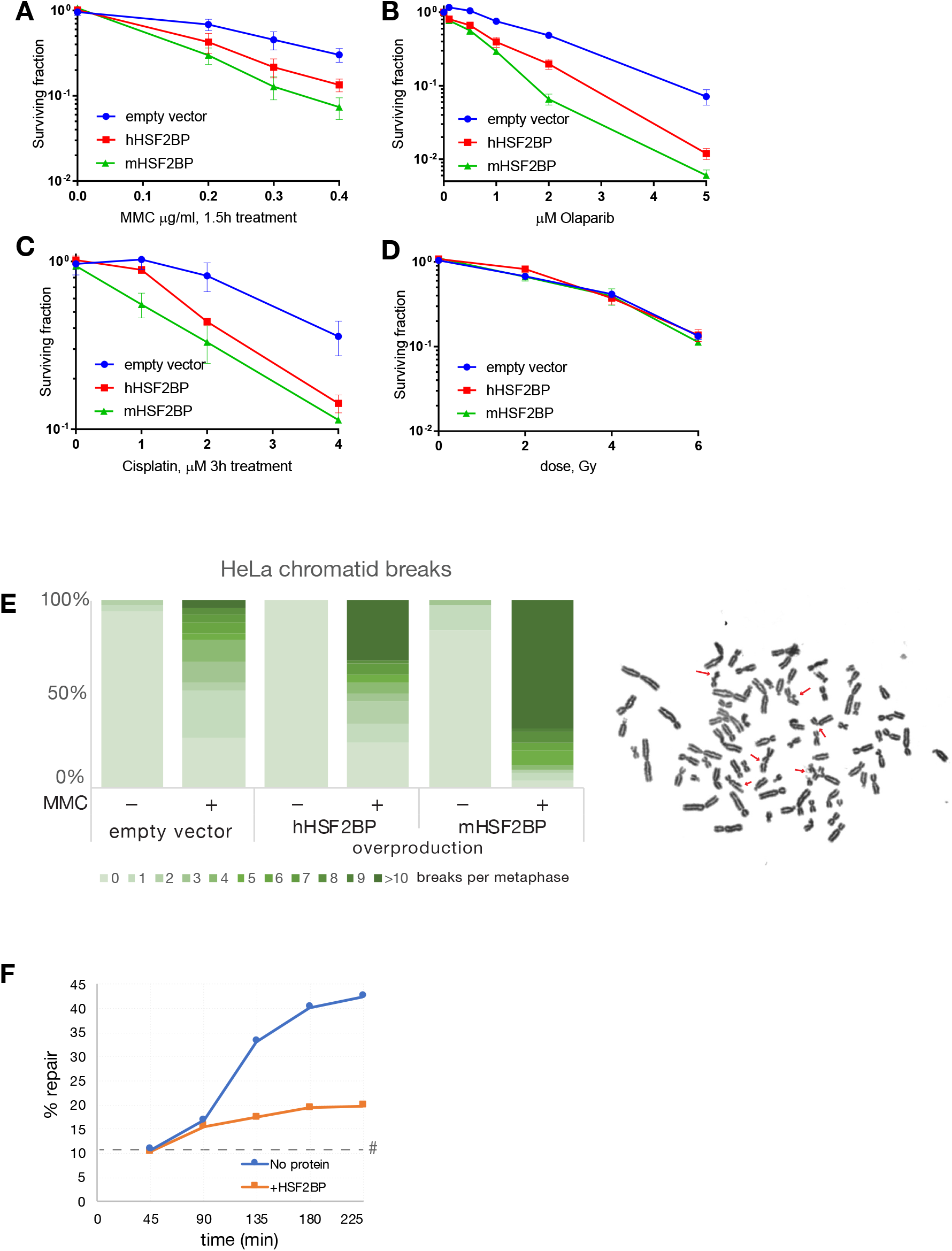
Overexpression of HSF2BP disrupts the Fanconi anemia pathway. HSF2BP overproduction disrupts the FA pathway (**A-D**) Clonogenic survivals of HeLa cells stably transformed with HSF2BP overexpression construct or empty vector. Each experiment was repeated at least two times with three technical replicates each. Error bars designate s.e.m. (**E**) Quantification of chromatid breaks in HeLa cells stably transformed with HSF2BP expression vectors or control, with and without overnight 100 nM MMC treatment. Example metaphase from HSF2BP overproducing cell treated with MMC is shown on the right, chromosomal aberrations indicated with arrows. (**F**) ICL repair **efficiency** in the presence or absence of HSF2BP. pICL was replicated in Xenopus egg extract that was supplemented with purified recombinant His-tagged hHSF2BP or buffer. Replication intermediates were isolated and digested with HincII, or HincII and SapI, and separated on agarose gel. Repair efficiency was calculated and plotted. As repair kinetics and absolute efficiency is highly dependent on the egg extract preparation single experiment is plotted here, and a replica is shown in **[Fig S2J]**. #, SapI fragments from contaminating uncrosslinked plasmid present in varying degrees in different pICL preparations. See also **[Fig S2K]**.

Hypersensitivity to DNA cross-linking agents in the absence of ionizing radiation sensitivity and proliferation defects is characteristic of cells deficient in FA proteins, rather than in general HR components. Efficient direct repeat GFP gene conversion in HSF2BP-overproducing cells **[Fig S2F]** indicated that HR was inherently functional, and normal cell cycle distribution **[Fig S2G]** suggested no replication defects. On the other hand, HSF2BP induced dramatic increase in chromosomal aberrations after treatment with MMC – a hallmark of FA cells **[Fig 2E]**. Consistent with the specific rather than generalized effect on BRCA2 function, BRCA2 concentration was not changed by HSF2BP overproduction **[Fig S2H]**, and although HSF2BP was predominantly cytoplasmic, BRCA2 remained nuclear **[Fig S2I]**. The last observation argues against cytoplasmic sequestration as the mechanism, as was previously proposed to explain the inhibitory effect of HSF2BP on transcription factor BNC1 (Wu et al., 2013).

To show that HSF2BP directly acts in the FA pathway of ICL repair, we took advantage of the evolutionary conservation of the HSF2BP-BRCA2 interaction **[Fig 1H]** and made use of the *Xenopus* egg extract-based ICL repair system. This system allows the repair of a single, site-specific cisplatin interstrand crosslink situated on a plasmid in a replication- and FA pathway-dependent manner (Knipscheer et al., 2012; 2009; Räschle et al., 2008). Using *Xenopus* egg extract, which does not contain endogenous xlHSF2BP at the level detectable by immunoblotting, we replicated the crosslinked plasmid (pICL) in the presence or absence of recombinant HSF2BP. Replication intermediates were isolated and repair efficiency was determined by measuring the regeneration of a SapI recognition site that is blocked by the ICL prior to repair. Addition of hHSF2BP almost completely abrogated ICL repair **[Fig 2F, S2J].** Importantly, addition of xlHSF2BP inhibited ICL repair to the same extent **[Fig S2K]**. These data suggest that ICL sensitivity of HSF2BP overexpressing cells is due to inhibition of ICL repair by the FA pathway.

### Mapping the BRCA2-HSF2BP interaction

To identify the regions required for the interaction between BRCA2 and HSF2BP we engineered a series of Flag-tagged BRCA2 fragment expression constructs and GFP-tagged HSF2BP expression constructs **[Fig 3A-C]**. For BRCA2, starting by dividing the protein into three fragments (N-terminal, middle, and C-terminal) **[Fig S3A]**, we mapped the HSF2BP-binding domain (HBD) to the 68 amino acid region (BRCA2-F9, Gly2270-Thr2337 in hBRCA2) located between the eighth BRC repeat (BRC8) and the DNA binding domain **[Fig 3C]**. Despite generally modest conservation of the region between BRC8 and the DNA binding domain, the BRCA2-F9 HBD is highly conserved and contains amino acid motifs that are perfectly conserved among vertebrate BRCA2 orthologues, including *X. laevis* **[Fig S3B]**. Deletion of the corresponding region from BRCA2 (BRCA2∆F9) essentially abolished the interaction **[Fig 3C, lane 6]**, therefore no other part of BRCA2 interacts with HSF2BP with high affinity.

**Figure 3.**
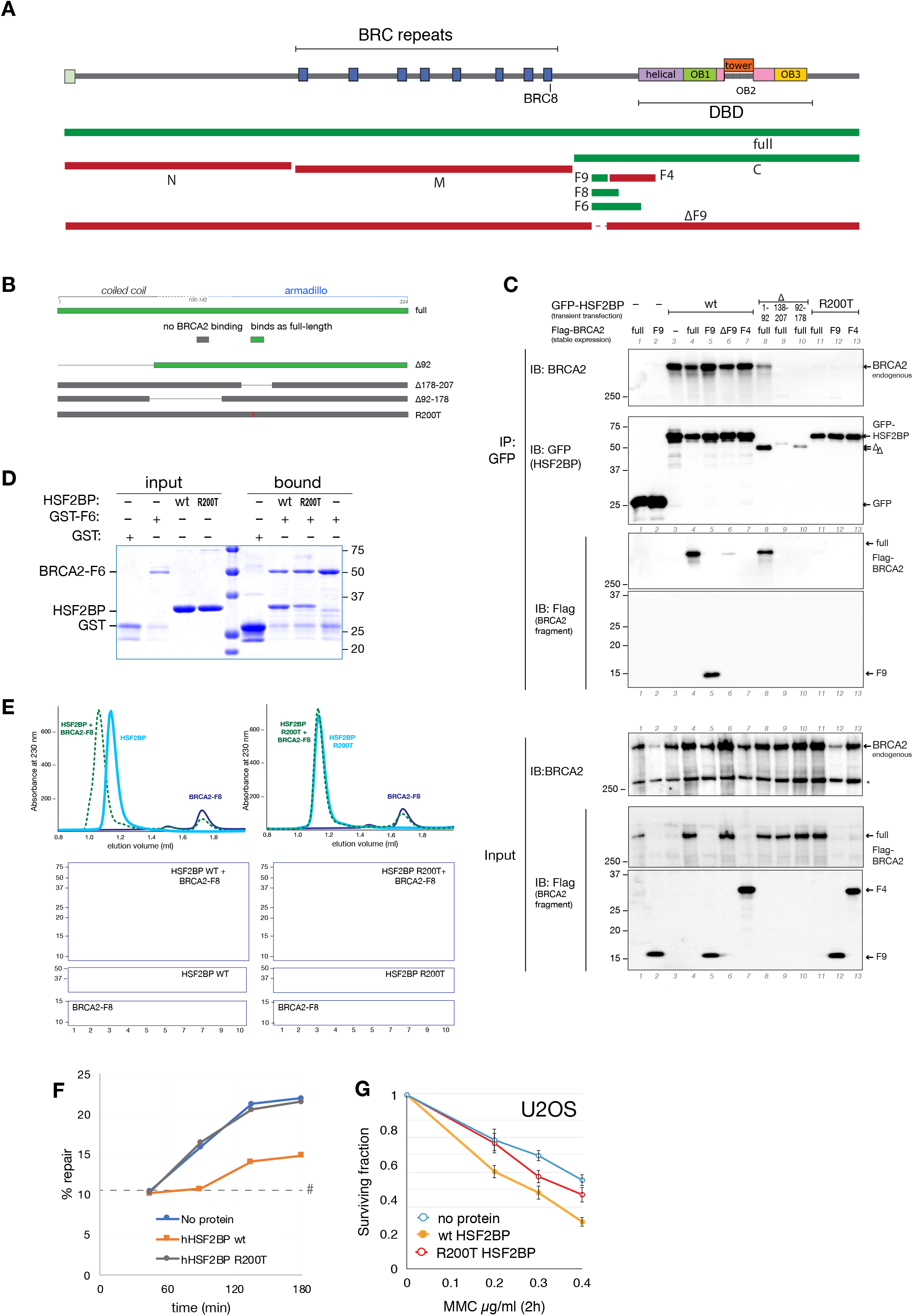
Mapping of BRCA2-HSF2BP interaction (**A**) Schematic of the BRCA2 protein domain structure, key BRCA2 fragments and internal deletion variant used to map the HSF2BP-BRCA2 interaction site. Red and green fills indicate, respectively, absence or presence of HSF2BP binding. (**B**) Schematic of the key HSF2BP variants used to map the residues involved in interaction with BRCA2. Boundaries between the putative coiled coil and the armadillo repeat regions are indicated. The armadillo repeat region is predicted starting at residues 100-140 with high confidence (e.g. 97% confidence for the model for A109-V334 of hHSF2BP by Phyre2 (Mezulis et al., 2015) based on β-catenin structure PDB 3SLA, which has 19% sequence identity). (**C**) Interaction between BRCA2 and HSF2BP revealed by co-immunoprecipitation between key variants. HeLa cells stably expressing Flag-tagged BRCA2 variants were transiently transfected with the indicated GFP-HSF2BP expression constructs (wt – full-length wildtype, ∆ – N-terminal and internal deletions, R200T point mutant) and anti-GFP immunoprecipitation was performed. (**D**) Direct interaction between purified recombinant HSF2BP (wild-type and R200T mutant) and GST-BRCA2 fragment F6 immobilized on glutathione sepharose beads. After washing bound proteins were released by heat denaturation, PAGE-fractionated and stained with CBB. (**E**) Interaction between purified untagged hHSF2BP and BRCA2-F8 fragment studied by analytical size exclusion chromatography. Proteins were analyzed separately (solid lines) or after co-incubation (dashed line). Complex formation between BRCA2-F8 and wild-type HSF2BP leads to an increase in hydrodynamic radius and elution in earlier fractions. This does not happen with R200T mutant. (**F**) HSF2BP R200T mutant does not inhibit ICL repair. pICL was replicated in Xenopus egg extract that was supplemented with recombinant His-tagged hHSF2BP WT or R200T mutant. Replication intermediates were isolated and digested with HincII, or HincII and SapI, and separated on agarose gel. Repair efficiency was calculated and plotted. A replica of this experiment is shown in **[Fig S3E]**. #, SapI fragments from contaminating uncrosslinked plasmid present in varying degrees in different pICL preparations. (**G**) HSF2BP R200T mutant does not sensitize human cells to MMC. Clonogenic survival of U2OS cells overproducing either wild-type or R200T mutant form of human HSF2BP (or transformed with empty vector) and treated with the indicated doses of MMC for 2 h one day after seeding. Data from three independent experiments is plotted, error bars indicated s.e.m.

Most of the GFP-fused truncated or internally deleted forms of HSF2BP we engineered were expressed at much lower levels than the full-length protein in human cells, presumably due to instability caused by misfolding **[Fig 3C, lanes 8-10]**. However, we could establish that deletions of the central and C-terminal parts of HSF2BP, which are predicted to adopt the armadillo repeat folds (e.g. Phyre2 (Mezulis et al., 2015)) abolished the interaction. Using internal deletions and eventually a set of seven point mutants **[Fig S3C]**, we established that this region is required for interaction with BRCA2, and that a single amino acid change in HSF2BP at position 200 from Arginine to Threonine (R200T) greatly reduces its ability to co-precipitate BRCA2 **[Fig 3C lanes 11-13]**. Notably, this residue is fully conserved among vertebrates. We could also establish that the N-terminal 92 amino acids, which are not predicted to contain armadillo folds and show weak sequence similarity to coiled coil regions, are not required for the interaction with BRCA2 **[Fig 3C lane 8]**.

To establish whether HSF2BP-BRCA2 interaction is direct, we purified bacterially expressed HSF2BP variants and BRCA2 fragments tagged with 6xHis or GST **[Fig 3D]** that could be cleaved off with TEV protease. Full-length HSF2BP was retained by GST-tagged BRCA2 F6 immobilized on glutathione sepharose beads, which demonstrated direct interaction between BRCA2 and HSF2BP. Consistent with the immunoprecipitation results, purified R200T mutant did not bind BRCA2-F6 efficiently **[Fig 3D]**. Analytical gel filtration of HSF2BP (wild-type or R200T) and BRCA2-F8, re-purified after cleaving off affinity tags, confirmed the formation of a complex between the two proteins **[Fig 3E]**. The hydrodynamic volume of HSF2BP was several fold higher than expected for a globular monomeric protein of its size (37.6 kDa), suggesting that HSF2BP exists in oligomeric and/or elongated form **[Fig S3D]**.

To show that the interaction with BRCA2 is important for the inhibition of the FA pathway by HSF2BP we monitored cisplatin ICL repair in *Xenopus* egg extract in the presence of wild type or R200T mutant hHSF2BP. While addition of wildtype HSF2BP strongly reduced ICL repair, addition of the R200T mutant protein showed comparable ICL repair efficiency to a condition where no protein was added **[Fig 3F, S3E]**. Consistent with this, overproduction of HSF2BP R200T did not significantly sensitize U2OS cells to MMC **[Fig 3G]**. This indicates that the HSF2BP-BRCA2 interaction is required for the inhibitory effect of HSF2BP on ICL repair.

### HSF2BP inhibits HR in the FA pathway

BRCA2 has a role in homologous recombination but also acts in the response to stalled replication forks. HSF2BP could therefore inhibit ICL repair at various stages of the repair process. Because ICLs are repaired synchronously in *Xenopus* egg extract we could investigate which repair step is affected by HSF2BP addition. An early key event in cisplatin ICL repair are the XPF-ERCC1-mediated nucleolytic incisions that unhook the ICL from one of the strands. These incisions can be directly observed during ICL repair in egg extract by replicating parentally labelled pICL **[Fig S4A]**. Replication intermediates are linearized and separated on denaturing agarose gels **[Fig S4B]**. The parental strand first migrates as a large X-shaped structure and during ICL unhooking it is converted to a linear molecule and arms (Klein Douwel et al., 2014; Knipscheer et al., 2009). We replicated prelabelled pICL in the presence and absence of hHSF2BP and measured the decline of the X-shape structures in time. No difference was observed in the presence or absence of HSF2BP indicating that ICL unhooking was not affected by the addition of HSF2BP **[Fig 4A]**.

**Figure 4.**
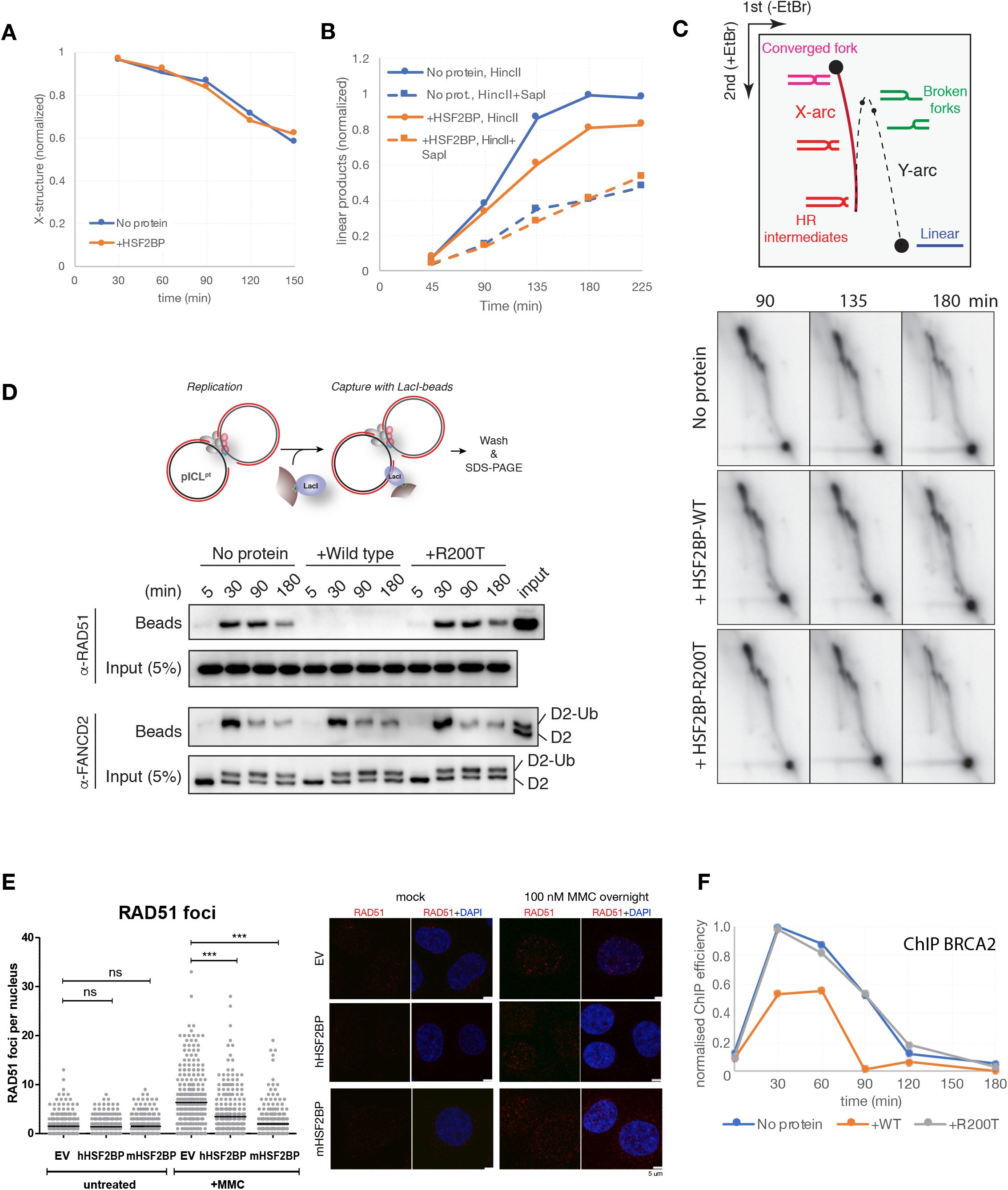
HSF2BP inhibits homologous recombination in the Fanconi anemia pathway. HSF2BP inhibits HR in the FA pathway (**A**) HSF2BP does not inhibit lesion unhooking in Xenopus egg extract. Prelabeled pICL was replicated in Xenopus egg extracts, replication products were isolated, digested by HincII, and separated on a denaturing agarose gel **[Fig S4B]**. The decline of the X-structures was quantified and plotted. See scheme in **[Fig S4A]**, main text and Methods for detailed description. (**B**) HSF2BP does not inhibit translesion synthesis in Xenopus egg extract. pICL was replicated in Xenopus egg extracts in the presence of 32P-lll-dCTP, replication products were isolated, digested by HincII, or HincII and SapI, and separated on a denaturing agarose gel **[Fig S4D]**. Extension products were quantified and plotted. Scheme of the assay in **[Fig S4C]**. (**C**) Inhibition of HR intermediate formation by HSF2BP revealed by 2D gel electrophoresis. pICL was replicated in Xenopus egg extracts in the presence of 32P- -dCTP, replication products were isolated, digested by HincII, and analyzed by 2D gel electrophoresis. (**D**) Stable RAD51 accumulation on pICL during repair is abrogated by HSF2BP wildtype, but not the R200T mutant. The pICL plasmid was pulled down by incubation with streptavidin beads coated with biotinylated LacI. Presence of bound RAD51 and FANCD2 was determined by immunoblotting. (**E**) Efficiency of RAD51 focus formation in HSF2BP-overproducing and control cells with or without 100 nM MMC treatment. After treatment cells were fixed, stained for immunofluorescence, mounted with DAPI and imaged using confocal microscopy. Images were scored manually for the number of foci. Data from three independent experiments is plotted, line represents average ± s.e.m., statistical significance was determined using the Kruskal-Wallis test with Dunn’s post-test. (**F**) HSF2BP inhibits BRCA2 loading at the ICL in Xenopus egg extract. pICL replication samples at the indicated time points were analyzed by BRCA2 ChIP.

After ICL unhooking a nucleotide is inserted across from the unhooked adduct by a currently unknown polymerase, followed by strand extension mediated by REV1 and polymerase ζ. This translesion synthesis (TLS) process can be monitored by separating nascently labelled pICL replication products that are linearized by digestion with HincII on a denaturing agarose gel. Replication stalling at the ICL site will initially generate arm fragments and these will be converted to fully extended linear fragments by TLS **[Fig S4C]**. Later, HR will result in the accumulation of additional full-length linear fragments that are cleavable by SapI. Of note, the TLS products are not cleavable by SapI because the unhooked adduct is not removed in *Xenopus* egg extract (Räschle et al., 2008). We separated HincII-digested nascent labelled repair products, that were replicated in the presence and absence of hHSF2BP, on a denaturing agarose gel and quantified the accumulation of linear products **[Fig S4D]**. While linear products accumulated in both conditions, the increase was less in the presence of HSF2BP **[Fig 4B]**. This could be caused by a defect in TLS and/or a defect in HR. To address this, we digested the repair intermediates with both HincII and SapI and performed the same analysis. The accumulation of full-length linear products was similar in absence or presence of HSF2BP **[Fig 4B]**. This shows that TLS is not affected by the addition of HSF2BP, and suggests that the reduction of linear products is caused by a defect in HR. To address this directly, repair intermediates were linearized and analyzed by two-dimensional gel electrophoresis (2DGE). In the absence of hHSF2BP, an X-arc was formed indicating the presence of HR intermediates (Long et al., 2011). In contrast, the presence of hHSF2BP but not its R200T mutant inhibited the formation of the X-arc **[Fig 4C, S4E]**, providing further evidence that HR is affected by HSF2BP.

### HSF2BP reduces loading of RAD51 and BRCA2 to ICLs

To address how HSF2BP inhibits HR during ICL repair we investigated BRCA2 and RAD51 recruitment to damage. To this end, we pulled down the pICL plasmid DNA during ICL repair in *Xenopus* egg extract using streptavidin beads coated with biotinylated LacI protein, which binds efficiently to nonspecific DNA (Budzowska et al., 2015). While this assay was not sensitive enough to detect BRCA2, RAD51 was readily accumulated on pICL in the absence of hHSF2BP **[Fig 4D]**. In contrast, no RAD51 was detected on pull downs in the presence of hHSF2BP **[Fig 4D]**. Consistent with this, RAD51 accumulation at sites of MMC-induced damage in human cells was significantly inhibited by HSF2BP **[Fig 4E]**. To investigate the binding of BRCA2 to the damaged DNA we then used chromatin immunoprecipitations (ChIP). BRCA2 was immunoprecipitated from ICL repair reactions in the presence and absence of HSF2BP, and the co-precipitated DNA was amplified by quantitative PCR with primers specific to the ICL region. BRCA2 accumulated at the ICL during repair as previously described (Long et al., 2014). In the presence of HSF2BP the accumulation of BRCA2 at the site of damage was reduced and the protein was unloaded earlier compared to conditions without added HSF2BP, or when the R200T variant of HSF2BP was added **[Fig 4F]**. The residual BRCA2 present at the ICL is most likely interacting with HSF2BP as we found accumulation of this protein by ChIP **[Fig S4F]**. In addition, ChIP experiments revealed normal recruitment of the unhooking endonuclease XPF and TLS polymerase REV1 in the presence of HSF2BP **[Fig S4G and H]** supporting our results that incisions and TLS are not affected. Moreover, FANCD2 recruitment and its ICL-dependent ubiquitination was normal both in *Xenopus* egg extract and in human cells **[Fig 4D, S4I and J]** indicating HSF2BP affects ICL repair downstream of FANCD2. Taken together these data demonstrate that HSF2BP affects the recruitment of both RAD51 and BRCA2 to the site of damage and thereby specifically inhibits HR during ICL repair.

## Discussion

Our study describes a novel, specific and evolutionarily highly conserved direct interaction between BRCA2, the key mediator of HR in vertebrates, and a previously functionally uncharacterized protein HSF2BP. We mapped the interaction to a short region within BRCA2, which despite its high sequence conservation had no described function. While a physiological role for HSF2BP may be restricted to germline, our work demonstrates the consequences of its ectopic activation. Using cell-biological methods in somatic human cells and biochemical assays in *Xenopus laevis* egg extracts we show that the presence of HSF2BP disrupts the HR step in the FA pathway of DNA ICL repair. Moreover, it sensitizes cells to ICL-inducing chemotherapeutic drugs and PARP inhibitors, and recapitulates the cytological phenotype characteristic of cells from FA patients.

While there is no indication that HSF2BP itself is involved in ICL or DSB repair (RNAi in **[Fig S2]**), analysis of ICL repair in *Xenopus* egg extract show that addition of HSF2BP specifically interferes with HR during this process. First, early ICL repair steps including FANCD2 recruitment, ICL unhooking, and translesion synthesis are not affected by HSF2BP, and neither is the recruitment of the factors involved in these steps. Second, the formation of HR intermediates is inhibited upon addition of HSF2BP. Third, RAD51 loading onto ICL containing plasmids is abrogated by HSF2BP. These data are also consistent with our finding that cells overproducing HSF2BP are sensitive to cisplatin and MMC, and show reduced RAD51 focus formation. How does HSF2BP affect HR? Several lines of evidence, including single particle tracking data, indicate that BRCA2 and HSF2BP form a stable complex in cells. ChIP analysis in *Xenopus* egg extracts indicate that this complex is recruited to ICLs during repair. In this complex, HSF2BP could affect RAD51 loading by a direct inhibition of BRCA2 function and/or an alteration in loading kinetics of BRCA2 onto the damage. In support of the latter, we showed reduced ICL recruitment and premature unloading of BRCA2 in the presence of HSF2BP. In fact, unloading precedes the formation of HR intermediates as observed on 2DGE. After 90 minutes no HR intermediates were visible but BRCA2 has already been completely unloaded in presence of HSF2BP. How HSF2BP promotes early unloading of BRCA2 is currently not known but this could involve regulated degradation or posttranslational modification. In addition, HSF2BP could inhibit BRCA2 function directly by preventing it from interacting with an important cofactor, or by affecting essential structural rearrangements. Why these mechanism specifically compromises HR during ICL repair, and not general HR, requires further investigation.

The sensitivity to ICL inducing agents and PARPi caused by ectopic HSF2BP expression, which is similar to FA protein deficiency, has an important implication, namely that the molecular mechanisms underlying FA-like genomic instability syndromes may include activation of wild-type genes. As FA (and phenotypically overlapping syndromes (van der Lelij et al., 2010)) are, with few exceptions (e.g. (Ameziane et al., 2015; Wang et al., 2015)), caused by deleterious recessive mutations affecting protein sequence, molecular diagnostics is mostly aimed at finding such mutations in genomic sequence. Elevated expression levels, which are central to cancer biology through the concept of oncogenes, are outside of the FA diagnostic paradigm. Yet there are examples of genetic diseases and developmental abnormalities caused by ectopic gene expression in humans and in model organisms (Epstein, 2009; Jaenisch and Bird, 2003; Lee and Young, 2013; Prelich, 2012). It is conceivable that *HSF2BP* ectopic (hyper)activation through mutation in *cis*-regulatory elements or transcription factors can arise, supported by the existence of *HSF2BP* overexpressing cancer cell lines and tumor samples. Although there may be confounding factors (see below), the finding warrants including gene expression analysis in the molecular characterization of patient cells from FA and clinically similar syndromes if conventional genetic testing fails to identify the cause.

The HSF2BP-binding domain (HBD) of hBRCA2 lies between the BRC8 repeat (~2051-2058) and the DNA-binding domain (~2484-3184). Two protein-protein interactions were mapped to this region: with DMC1, the meiosis-specific paralogue of RAD51 (via the PhePP motif in 2386-2411 of hBRCA2 (Thorslund et al., 2007)), and with FANCD2 (hBRCA2 2350-2545 (Hussain et al., 2004)), however neither overlaps the HBD (2270-2337).

Based on HSF2BP expression profile and the phenotypes we described, it does not seem likely that mutations in the HBD would predispose to cancer by disrupting the HSF2BP-BRCA2 interaction. On the contrary, such mutation would be protective against genomic instability induced by HSF2BP. On the other hand, since we show that HSF2BP binding to HBD is required for ICL repair disruption, it is plausible that HBD itself is involved in a molecular transaction required for ICL repair, which is blocked by HSF2BP. In this case, missense mutations in HBD could be pathogenic. We surveyed several curated databases combining clinical, functional and sequence data on *BRCA2* (Béroud et al., 2016; Fokkema et al., 2011; Szabo et al., 2000) and found two experimentally characterized variants (R2336H and I2285V). The R2336H variant is classified as potentially deleterious or causal, based on cellular HR assay, but its effect is attributed to aberrant splicing and BRCA2 protein truncation (Biswas et al., 2011; Claes et al., 2004). The position is not well conserved evolutionarily **[Fig S3B]**. The second missense variant p.I2285V (c.6853A>G) is associated with increased frequency of exon 12 skipping (Bièche and Lidereau, 1999; Li et al., 2009; Rauh-Adelmann et al., 2000). Exon 12 encodes half of the F9 fragment **[Fig S3B]** and can be spliced out without a reading frame shift. The *BRCA2* ∆12 transcript variant is present natively in a range of cancer lines and breast tumor tissues. Extensive characterization of mES cells expressing only ∆12 transcript revealed no detectable phenotype (Li et al., 2009) and no breast cancer predisposition was found among c.6853A>G carriers, which led to the conclusion that increase in ∆12 spice variant in breast cancer cell lines is an epiphenomenon (Li et al., 2009). However, it is tempting to speculate, based on our findings, that *HSF2BP* overexpression followed by shift to BRCA2∆12 could be beneficial for cancer cell evolution, by first inducing genomic instability leading to transformation, and then reverting back to the more efficient DNA repair for robust proliferation. The ratio between BRCA2 splice variants may explain potential differences in sensitization due to HSF2BP overproduction in different cell lines. Given high conservation of the region among vertebrates, including near complete conservation of I2285 and the surrounding positions, it is possible that homozygous loss of exon 12 does have negative consequences in meiosis, which could not have been detected in the published functional and genetic analyses (Li et al., 2009).

Overproduction of HSF2BP sensitized cells to PARPi to a similar extent as to ICL agents. Several reports noted increased sensitivity of FA cells to PARPi: a panel of isogenic chicken DT40 cell mutants (Murai et al., 2012), fibroblast from FA mouse models (McCabe et al., 2006), human and mouse head and neck cancer lines (Lombardi et al., 2015), patient lymphoblastoid lines (Stoepker et al., 2015). However, PARPi sensitivity of core FA proteins and I/D2 is much lower than in BRCA2-deficient cells, and lack of sensitivity was concluded for FA-A, -L, -D2, -I, -J patient lines(Kim et al., 2013). To the extent a comparison between disparate cell lines can be made, sensitization by HSF2BP overproduction is closer to PARPi sensitivity of core FA and FA-D2/I cells than to the much more severe sensitivity of cells deficient in general HR (BRCA2, PALB2, RAD51, BRCA1) or FANCP/SLX4. This is consistent with an inhibitory role for HSF2BP in HR during ICL repair but not in general HR.

Our data demonstrate that ectopic (over)expression of *HSF2BP* can cause genomic instability specifically due to inefficient ICL repair, while other HR reactions remain unaffected: cells proliferate normally and the steps preceding HR intermediate formation in *Xenopus* egg extract, all of which are replication-dependent, are also unaffected. This specificity is important for two reasons. First, it is emerging that accumulation of genomic instability due to gradual segmental loss of the *BRCA2* function is likely to be more conducive to oncogenic transformation than the catastrophic consequences of the complete inactivation of this essential gene (Feng and Jasin, 2018; Hartford et al., 2016; Tan et al., 2017). Second, we anticipate that the separation of general and ICL-specific HR functions of BRCA2, which we could achieve in cultured cells and in biochemical assays, will be as useful for understanding the complex biology of BRCA2 as was the separation of its DSB repair and of replication fork stabilization functions (Lomonosov et al., 2003; Schlacher et al., 2012). Finally, this novel mechanism of ICL repair inhibition could be used to for the development of more efficient treatment for cancer patients with interstrand crosslinking agents.

## Acknowledgements

The authors would like to thank Dr John Martens for providing breast cancer lines HSF2BP and for help with analysis of expression data, G.Stier from EMBL for the kind gift of expression vectors pETM11 and pETM30. KS was supported by the Uehara Memorial Foundation, the Mochida Memorial Foundation for Medical and Pharmaceutical Research, and the JSPS Postdoctoral Fellowship for Research Abroad. PK was supported by the Netherlands organization for Scientific Research (VIDI 700.10.421) and a project grant from the Dutch Cancer Society (KWF HUBR 2015-7736). J.L is supported by the gravitation program from the Netherlands Organization for Scientific Research (NWO). The research leading to these results has received funding from the European Community’s Seventh Framework Program (FP7/2007-2013) under grant agreement No. HEALTH-F2-2010-259893 and from the Dutch Cancer Society (grant EMCR 2008-4045 and a Ride for the Roses Cancer Research Grant).

## Author Contributions

AZ and RK conceived the study. IB, AZ, SvR-F, MR, HO, NV, NvdT, AO performed human cell and biochemical experiments. KS performed Xenopus egg extract experiments. DD and KB collected mass spectrometry data. JCD, JD, JL, CW, DvG, PK, AZ and RK supervised different parts of the study. AZ, PK, RK and KS wrote the manuscript with contributions from other authors. All authors read and approved the manuscript.

## Declaration of Interests

The authors declare no competing interests.

## Methods

### Cell culture

HeLa, U2OS, MDA-MB-157 and RT-112 cells were cultured in DMEM, supplemented with 10 %(v/v) FCS and penicillin/streptomycin. HCC1187 were cultured in RPMI-1640 supplemented with 10% FCS and penicillin/streptomycin. Gene-targeted mouse ES cells were derived from the IB10 cell line, which is a subclone of E14 129/Ola from male origin, specific pathogen free, were cultured on gelatinized plastic dishes (0.1% gelatin in water) as described before (Zelensky et al., 2017) in media comprising 1:1 mixture of DMEM (Lonza BioWhittaker Cat. BE12-604F/U1, with Ultraglutamine 1, 4.5 g/l Glucose) and BRL-conditioned DMEM, supplemented with 1000 U/ml leukemia inhibitory factor, 10% FCS, 1x NEAA, 200 U/ml penicillin, 200 μg/ml streptomycin, 89 μM β-mercaptoethanol.

### Generation of genetically modified cell lines

*Hsf2bp^wt/GFP^* mES cells were created by gene targeting using the approach we used previously to engineer the BRCA2-GFP knock-in lines (Reuter et al., 2014). The gene targeting construct was engineered by recombineering, starting with the BAC clone bMQ-430H02 from the Sanger 129/Sv library 43, homology arms were 3.1 and 6.2 kb long, boundaries defined by the retrieval primers HSF2BP-CG-rtrL and HSF2BP-CG-rtrR, and the GFP-2A-neo cassette 19 was inserted after the last codon of the Hsf2bp CDS, using the primers HSF2BP-CG-F and HSF2BP-CG-R. After the completion of selection with 200 μg/ml G418 clones were isolated. Correct targeting was confirmed by Southern blotting on HindIII-digested genomic DNA with a probe produced by PCR amplification with the primers HSF2BP-pr5-F and HSF2BP-pr5-R. Two out of the 14 screened clones were identified as correctly recombined. *Rad51ap1^wt/GFP^* cells used as a control in the mass spectrometry experiment were produced by similar gene targeting procedure (BAC bMQ-338B20, 3.6 and 4.8 kb homology arms) and converted to homozygosity by high-G418 selection as described before (Reuter et al., 2014). The gene targeting construct for the production of *Palb2^GFP/GFP^* cells was engineered using the same approach (BAC bMQ-128G09, 3.7 kb and 4.4 kb right homology arms), but *S. pyogenes* CRISPR/Cas9 was used to stimulate recombination. The target for sgRNA cloned into pX459 vector (Ran et al., 2013) was 5’-tatataccgatacttttaag-3’. With the exception of U2OS GFP-HSF2BP clone #5, all stable cell lines were constructed using PiggyBac vectors by co-transfecting them with the transposase expression construct (mPB or hyPBase), followed by selection with either 1.5 μg/ml puromycin or 800 μg/ml G418 was started and maintained for 6-10 days. Stable transformation was highly efficient (>95% GPF+ cells when GFP-tagged constructs were used) and therefore clonal isolation was not performed. Stable line U2OS GFP-HSF2BP clone #5 used in the initial experiment was constructed by random integration of pEGPFN1-HSF2BP construct with 800 μg/ml G418 selection. Several GFP+ individual clones were isolated, expanded and checked for uniform expression of single major GPF-fused protein by FACS and immunoblotting.

### Mass spectrometry

Stable isotope labeling in cell culture (SILAC) of mES cells was performed by expanding the cell lines from 6 cm to 2x 145mm dishes (gelatinized) in drop-out ES media (Mouse Stem Cell Expansion DMEM for SILAC, (#88207 Thermo Fisher) to which 3.5 mg/ml D-glucose, 105 μg/ml L-leucine, 50 ml dialyzed FCS, 6 ml ultraglutamine (Lonza BE17-605E/U1) were added) complemented with 84 μg/ml L-lysine (K) and 146 μg/ml L-arginine (R). Lysine and arginine differed in N and C isotopic content for three SILAC states: K0R0 for light, K3R6 for medium and K8R10 for “heavy”; heavy-isotope amino acids were purchased from Cambridge Isotope Laboratories. Sample preparation after pull-down with anti-GFP nanobody agarose beads (Chromotek) involved either PAGE fractionation or on-beads digestion, followed by HPLC. Data acquisition was performed using LTQ-Oribtrap XL instrument (Thermo) and analyzed using MaxQuant software (Cox et al., 2009).

### Expression Constructs

The CDS of human and mouse HSF2BP were PCR-amplified from first-strand cDNA synthesized from total RNA isolated from U2OS and mES IB10 cells, respectively, with SuperScript II enzyme (Invitrogen) and oligo-dT primer. The PCR products were cloned into pCR4 topo-TA vector (Invitrogen) and verified by Sanger sequencing. Clones containing sequences corresponding to the predicted full-length HSF2BP CDS as annotated the NCBI GenBank database (NM_007031 and NM_028902) were used as templates for subcloning. Re-cloning into destination vector was performed using isothermal Gibson assembly (Gibson et al., 2009). The GFP expression constructs used in the initial experiments were derived from pEGFP-N1 and pEGFP-C1 vectors for C- and N-terminal fusions, respectively. Other GFP- and Flag-tagged expression constructs were assembled in PiggyBac described before (pAZ125 (Zelensky et al., 2017)) or engineered using analogous steps in our lab. The PiggyBac vectors carried PGK-neo or PGK-puro selection cassettes and CAG promotor-driven transgene as a separate expression unit. Human BRCA2 CDS fragment were PCR-amplified or excised using restriction from phCMV-MBPx2-BRCA2 vector (Jensen et al., 2010) and cloned. All cloning junctions and fragments produced by PCR were sequence-verified after cloning. Internal deletions were produced by excising a fragment of full-length Flag-BRCA2 PiggyBac expression construct (pAZ114) using two unique restriction sites nearest to the region targeted for deletion and patching the gap with appropriate PCR-amplified fragment(s) encoding the desired deletion using Gibson assembly. Site-directed mutagenesis of GFP-HSF2BP was performed by amplifying two overlapping fragments with the mutation encoded by the overlapping part of the primers (20-25 bp) and cloning the two fragments into destination GFP PiggyBac vector. Proofreading Q5 PCR polymerase (NEB) was used and clones were verified by Sanger sequencing. Details of the PCR primers and cloning strategies are available upon request.

### Lentiviral shRNA transduction

shRNA expression constructs were obtained from the Sigma mission library (TRC 1.5). Lentiviral packaging plasmids (pMDLg/pRRE, pRSV-REV and pMD2.G) and shRNA expression constructs were transfected into HEK293T cells using calcium phosphate precipitation. 24h after transfection the medium was changed and 48 h after transfection the supernatant of the HEK293T cells was added to U2OS cells. This process was repeated the next day. Forty-eight hours after the second transduction, selection with 1.5 μg/ml puromycin was started. Two of the five tested shRNAs efficiently reduced the concentration of overproduced GFP-HSF2BP: #2 (clone TRCN0000017473, target sequence GCTGGAATTGTCACGAATGTT) and #3 (clone TRCN0000017476, target sequence GCTAATGCTGATGTCCCTATA); constructs SHC001 (empty vector) and SHC003 (non-targeting shRNA) from the same library were used as negative controls.

### FACS-based techniques

Two parameter cell cycle analysis was performed as described (Zelensky et al., 2013) but a BD LSRFortessa cell analyzer was used. For DR-GFP assay, U2OS cells carrying the reporter (clone U2OS-18 (Puget et al., 2005)) were transfected with the PiggyBac expression construct for HSF2BP or empty vector, and the construct encoding transposase, stable integrants were selected with 800 μg/ml G418. The stably transformed U2OS-18 derivatives were seeded into 6-well plates at 2*10^5^ cells per well and transfected next day using XtremeGene reagent with 2 μg of either I-SceI expression plasmid (pCBASceI), empty vector, or pEGFP-N1 as transfection control efficiency control. FACS analysis was performed two days later.

### Antibodies

Antibodies used in this study were against RAD51 (rabbit 2307 (Tan et al., 1999)), FANCD2 (NB 100-316, Novus Biologicals), H2B (07-371, Millipore), MSH2 (Ab-2, Oncogene), HSP90 (ab13492, Abcam), BRCA2 (Ab-1, OP95, Calbiochem), FLAG (M2 antibody, Sigma), GFP (clones 7.1 and 13.1, Roche), ORC2 (551178,BD Pharmingen), PARP-1 (C2-10, Enzo), 6xHis tag (ab18184, Abcam), and XRCC3 (ab6494, Abcam). Anti-HSF2BP rabbit polyclonal antibodies SY8126 and SY8127 were raised against purified recombinant untagged human HSF2BP (Kaneka Eurogentec, Belgium) and used either as crude serum or after affinity purification against GST-HSF2BP immobilized on glutathione sepharose as described (Chalkley and Verrijzer, 2004). Antibodies against xlREV1, xlXPF, and xlFANCD2 were previously described (Budzowska et al., 2015; Klein Douwel et al., 2014; Räschle et al., 2008; Walter & Newport, 2000). The BRCA2 antibody was raised against residues 1842-2080 of xlBRCA2. The cDNA encoding the fragment was codon-optimized for *E. coli*, synthesized (gBlocks Gene Fragments, Integrated DNA Technologies), and ligated into the XhoI-BamHI sites of the pETDuet-1 vector (Novagen). The fragment was overexpressed in *E. coli* BL21(DE3) cells (New England BioLabs) with a N-terminal His-tag, and purified by the method described previously (Klein Douwel et al., 2014). The purified antigen was used for immunization of rabbits (PRF&L, Canadensis, USA). Specificity of the antisera was confirmed using western blotting.

### Immunoblotting and Cell fractionation

To prepare total protein lysates, cells were scraped in PBS and lysed in 2x Laemmli SDS loading buffer (120 mM Tris pH 6.8, 4% SDS, 10% Glycerol), after determining the protein concentration in the lysate by Lowry method, the lysate was complemented by 10x reducing additive (0.1% bromophenol blue, 0.5% β-mercaptoethanol). For fractionation 1 million cells were collected by trypsinization, washed with ice-cold PBS, re-suspended in Schaffner’s lysis buffer (10 mM HEPES-NaOH pH 7.9, 10 mM KCl, 0.1 mM EDTA, 0.1 mM EGTA, 1 mM DTT, protease inhibitor cocktail); after 15-min incubation on ice NP-40 was added to the final concentration 0.1% and solution was vortexed. Nuclei were pelleted by 30 sec centrifugation at 18,000 rcf, re-suspended by shaking for 15 min at 4 °C in nuclear extraction buffer (20 mM HEPES-NaOH pH 7.9, 400 mM KCl, 1 mM EDTA, 1 mM EGTA, 1 mM DTT, protease inhibitor cocktail) and centrifuged for 5 min at 18,000 rcf to separate chromatin pellet from soluble nucleoplasmic fraction; the pellet was re-suspended in Laemmli SDS loading buffer. Proteins were separated on polyacrylamide handcast tris-glycine, or precast bis-tris or tris-acetate gels (Novex) and blotted on nitrocellulose or PVDF. For BRCA2 detection 4-8% precast tris-acetate gels were used, and transfer was performed in 2x Towbin transfer buffer (50 mM Tris, 384 mM glycine, 20% methanol) at 300 mA constant current for 2h at 4 °C to PVDF membrane. Membranes were blocked in 3% milk in PBS+0.05% Tween. After overnight incubation with the primary antibody, membranes were washed in PBS+0.05% Tween and incubated with horseradish peroxidase-conjugated secondary antibodies (Jackson Immunoreserach). Blots were developed using homemade ECL reagents and detected with the Alliance 4.7 (UVItec).

### Protein Purification

Human HSF2BP cDNA was re-cloned into pETM11 (6xHis-TEV) expression vector using Gibson assembly and transformed into *E. coli* Rosetta2 (DE3) pLysS. A starter culture (LB + 50 μg/ml kanamycin, 30 μg/ml chloramphenicol) was grown overnight at 37 °C, used to inoculate 4.5 L of LB which was grown at 37 °C until the OD_600_ reached 0.6. Expression was induced by addition of IPTG to 0.2 mM, incubation was continued at 16 °C for 16 h, cells were harvested and frozen, thawed in equal volume of 2x lysis buffer (1M NaCl, 25mM Tris pH7.5, 5% Glycerol, 5 mM β-mercaptoethanol, 1x EDTA-free protein inhibitor cocktail (Roche)) and sonicated (8x 30 sec). The lysate was cleared by centrifugation at 35,000 rcf for 45 min, and loaded on Ni-NTA beads equilibrated with binding buffer (500 mM NaCl, 25mM Tris pH7.5, 5% Glycerol, 5 mM β-mercaptoethanol). Beads were washed with increasing concentration of imidazole (20 and 40 mM) in binding buffer. Fractions were eluted with 250 mM imidazoleand analyzed by SDS-PAGE. Fractions containing HSF2BP were pooled and loaded on Superdex 200 16/60 gel filtration column equilibrated with GF buffer(250 mM NaCl, 25 mM Tris pH7.5, 5% Glycerol, 5 mM β-mercaptoethanol) on an ÄKTA FPLC system (GE Healthcare). Peak fractions containing HSF2BP were pooled and further purified on a 5 ml Hitrap Q anion exchange column using a gradient from 100 till 600 mM NaCl in 25 mM Tris pH7.5, 5% Glycerol, 5 mM β-mercaptoethanol. Fractions containing HSF2BP were pooled, concentrated, flash-frozen in GF buffer using liquid nitrogen and stored at -80°C. To produce HSF2BP without his-tag, HSF2BP eluted from the Ni-NTA beads was mixed with Tobacco Etch Virus (TEV) protease and dialyzed o/n at 4°C against GF buffer before loading onto the gel filtration column. HSF2BP variant R200T was purified as wild type. HSF2BP concentration was determined spectrophotometrically (ϵ^280nm^ = 20970 M^-1^ cm^-1^).

BRCA2 fragment F6 was cloned into pETM-30 (6xHis-GST-TEV) expression vector and purified and stored according to the protocol described above for HSF2BP, without removal of the tag. Concentration of the fusion protein was determined spectrophotometrically using ϵ^280nm^ = 52830 M^-1^ cm^-1^. BRCA2 fragment F8 was cloned into pETM11 and purified and stored according to a protocol similar to that for HSF2BP,up to and including TEV cleavage to remove the tag. Subsequently the dialysed material was loaded onto a 5 ml Hitrap S cation exchange column. BRCA F8 eluted in the flow through, was concentrated and further purified on a Superdex 75 16/60 size exclusion chromatography column (GE Healthcare) equilibrated with GF buffer. Concentrated fractions were flash frozen and stored at -80°C. The F8 concentration was estimated using SDS-PAGE with Coomassie Brilliant Blue staining, using bovine serum albumin (BSA) as the standard.

The cDNA encoding full-length xlHSF2BP was codon-optimized for *E. coli*, synthesized (gBlocks Gene Fragments, Integrated DNA Technologies), and ligated into XhoI-BamHI sites of the pETDuet-1 vector. xlHSF2BP was overexpressed with a N-terminal His-tag in *E. coli* BL21(DE3) cells at 18 °C. The cells were collected by centrifugation, resuspended in buffer A (50 mM Tris pH 8.0, 10% glycerol, 500 mM NaCl, 1 mM phenylmethylsulphonyl fluoride, 10 mM imidazole, and 5 mM DTT), and disrupted by sonication. The soluble fraction was collected by centrifugation at ~40,000 × g for 25 min at 4 °C, and mixed gently with 1 ml Ni-NTA agarose resin (Life Technologies) at 4 °C for 1 h. The Ni-NTA agarose resin was packed into Poly-Prep chromatography column (Bio-Rad), and washed with 50 ml buffer A containing 20 mM imidazole. The xlHSF2BP protein was eluted with 8 ml buffer A containing 400 mM imidazole, and concentrated using a 30 kDa MWCO Amicon Ultra-15 centrifugal filter unit (Millipore). The sample was then loaded onto a Superdex 200 column (HiLoad 16/60 preparation grade, GE Healthcare) equilibrated with buffer B (25 mM Tris pH 7.5, 5% glycerol, 200 mM NaCl, and 5 mM DTT). The eluted xlHSF2BP protein was concentrated, and aliquots were flash frozen with liquid nitrogen. The protein concentration was determined by SDS-PAGE with Coomassie Brilliant Blue staining, using bovine serum albumin (BSA) as the standard protein.

### Analytical Gel Filtration

A Superdex 200 increase 3.2/300 (GE Lifesciences) was equilibrated with GF buffer on an ÄKTA Micro system (GE Healthcare). In a total volume of 50μl, 37μM untagged HSF2BP (wild type or R200T) and/or 60μM BRCA2-F8, was applied to the column at a flow of 50 μl/min. 50 μl fractions were collected and analyzed using SDS-PAGE (15 % acrylamide).

### GST Pull-down

20μl of glutathione beads were prewashed with buffer (150 mM NaCl, 25mM Tris pH7,5, 5% Glycerol, 5mM β-mercaptoethanol). 200 μmol of GST or GST-BRCA2 F6 was bound for 1 h at 4^<u>o</u>^ C. Beads were washed 3x with buffer. 500 μmol of HSF2BP (WT or R200T) was added and incubated for 1hr at 4C. Beads were washed 3x with buffer. Samples were analyzed on SDS-PAGE.

### *Xenopus* egg extracts, DNA replication, repair assay, and lesion bypass assay

DNA replication and preparation of *Xenopus* egg extracts (HSS and NPE) were performed as described previously (Tutter and Walter, 2006; Walter et al., 1998). Preparation of plasmid with a site-specific cisplatin ICL (pICL), and ICL repair assays were performed as described (Enoiu et al., 2012; Räschle et al., 2008). Briefly, pICL (9 ng/μL) and pQuant (0.45 ng/μL) were first incubated in a high-speed supernatant (HSS) of egg cytoplasm for 20 min at ~ 20 °C, which promotes the assembly of prereplication complexes on the DNA. Addition of two volumes nucleoplasmic egg extract (NPE), which also contained 32P-α-dCTP, triggers a single round of DNA replication. Where indicated, his-tagged hHSF2BP (0.45 μM) or his-tagged xlHSF2BP (0.45 μM) was added to NPE prior to mixing with HSS. Aliquots of replication reaction (3.8 μl) were stopped at various times with 45 μl Stop solution II (50 mM Tris pH 7.5, 0.5% SDS, and 10 mM EDTA,). Samples were incubated with RNase (0.13 μg/μl) for 30 min at 37 °C followed by proteinase K (0.5 μg/μL) overnight at room temperature. DNA was extracted using phenol/chloroform, ethanol-precipitated in the presence of glycogen (30 mg/ml), and resuspended in 3.8 μl TE (10 mM Tris pH 7.5 and 1 mM EDTA). ICL repair was analyzed by digesting 1 μl extracted DNA with HincII, or HincII and SapI, separation on a 0.8% agarose gel in 1x TBE buffer, and quantification using Typhoon TRIO+ (GE Healthcare) and ImageQuant TL software (GE Healthcare). Repair efficiency was calculated as described (Knipscheer et al., 2012). For lesion bypass assay, the digested DNA was ethanol-precipitated, and resuspended in 12 μL alkaline loading buffer (50 mM NaOH, 1 mM EDTA, and 2.5 % Ficoll-400). Fragments were then separated on a 0.8% agarose gel in alkaline buffer (50 mM NaOH and 1 mM EDTA), after which the gel was dried on Amersham Hybond-XL membrane (GE Healthcare) and exposed to a phosphor screen. The band intensity of the bypassed product was quantified with Typhoon TRIO+ using ImageQuant TL software. The highest value was set at 100% for the bypassed product.

### Immunoprecipitation (IP)

For co-IP mES cells were washed twice in ice-cold PBS and lysed in situ in NETT buffer (100 mM NaCl, 50 mM Tris pH 7.5, 5 mM EDTA pH 8.0, 0.5% Triton-X100) supplemented immediately before use with protease inhibitor cocktail (Roche) and 0.4 mg/ml Pefabloc (Roche) (NETT++). Four hundred fifty μl NETT++ buffer was used per 145 mm dish. After 30 min lysis on ice, mixtures were centrifuged and the supernatant (input) was added to washed anti-GFP beads (Chromotek) or anti-Flag M2 affinity gel (A2220, Sigma)). Beads and lysates were incubated 2-4 h at 4 °C while rotating, washed three times in NETT++ buffer and bound proteins were eluted by boiling in 2x Sample buffer. Flag IPs were performed using M2 agarose gel (Sigma, A2220). For proteins smaller than 75 kDa Glycine elution according to the manufacturers protocol was used to prevent release of the M2 antibody from the affinity gel.

IP from *Xenopus* egg extracts was performed as described previously with modifications (Knipscheer et al., 2009). 3 μl BRCA2 or HSF2BP antiserum was incubated with 30 μl Protein A Sepharose Fast Flow (PAS) beads (GE Healthcare) overnight at 4 °C. The beads were washed twice with 400 μl PBS, once with 400 μl ELB (10 mM HEPES–KOH (pH 7.7), 50 mM KCl, 2.5 mM MgCl2, and 250 mM sucrose), twice with 400 μl ELB supplemented with 0.5 M NaCl, and finally twice with 400 μl ELB supplemented with 0.25 mg/ml BSA. For IP with BRCA2 antiserum, 5 μl of the antibody-bound beads were incubated with 20 μl diluted NPE (4 μl NPE and 16 μl ELB) for 1 hl at 4 °C. 1.5 μl HSF2BP (3 μM) was then added to the beads, and the reaction mixture was incubated for 3 h at 4 °C. The beads were washed 4 times with 400 μl ELB containing 80 mM NaCl and 0.5% Triton-X100 (Sigma). For IP with HSF2BP antiserum, 5 μl of the antibody-bound beads were incubated with 2 μl HSF2BP (3 μM) for 1 h at 4 °C, and washed twice with 400 μl ELB supplemented with 0.25 mg/ml BSA. The beads were then incubated with 20 μl diluted NPE (4 μl NPE and 16 μl ELB) for 3 h at 4 °C, and washed 4 times with 400 μl ELB. The bound proteins were eluted by adding 10 μl 1x SDS sample buffer (75 mM Tris pH 6.8, 10% glycerol, 2.5% SDS, 50 mM TCEP (Bond-Breaker TCEP solution, Life Technologies), and 0.025 % Bromophenol blue), separated by SDS-PAGE, and detected by western blotting with RAD51 (1:10,000), BRCA2 (1:1,000), or HSF2BP (1:2,500) antiserum. Preimmune serum was used for mock immunoprecipitation.

### Immunofluorescence and Microscopy

Direct HSF2BP-GFP imaging and immunofluorescence staining was performed on ES cells grown overnight on a glass coverslip coated with laminin, which improves their attachment and morphology. Sterile 24 mm coverslip was placed in a 6-well plate, and a 100 μl drop of 0.05 mg/ml solution of laminin (Roche, 11243217001) was pipetted in the middle of it. The plate was left for ~30 min in the cell culture incubation, after which the laminin solution was aspirated and cell suspension was placed in the well. Cells were fixed for 15 min in 2% paraformaldehyde in PBS at room temperature and either mounted directly in VectaShield with DAPI or immunostained with anti-53BP1 antibody.

Oblique illumination microscopy and particle tracking were performed exactly as described (Reuter et al., 2014). The experiment was repeated three times, two independent *Hsf2bp^GFP/+^* clones were used in each.

HeLa and U2OS cells were treated overnight with 100 nM MMC, pre-extracted before fixation for 1 min in Triton X-100 buffer (0.5% Triton X-100, 20 mM HEPES-KOH pH 7.9, 50 mM NaCl, 3 mM MgCl2, 300 mM sucrose), fixed in 2% paraformaldehyde in PBS and stained as described (Modesti et al., 2007). For quantification, experiments were performed in triplicate and per sample 100 nuclei were counted. Pictures were taken using a Leica SP5 confocal microscope using a 63x objective and 405, 488 and 594nm lasers. Statistical analysis was performed in Graphpad Prism using a Kruskal-Wallis test with Dunn’s post-test.

### Incision assay

Incision assay was performed as described previously (Klein Douwel et al., 2014; 2017). Briefly, pICL (225 ng) and pQuant (11.3 ng) were incubated with 1.5 units NB-BSR DI enzyme (NEB) in 1× NEBuffer 2.1 for 30 min at room temperature. Subsequently, the nick translation reaction was initiated by adding 11 μl DNA Polymerase I mix (5 units DNA polymerase I (NEB), dATP, dGTP, dTTP (0.5 mM each), dCTP (0.4 μM), ^32^P-α-dCTP (3.3 μM) in 1× NEBuffer 2.1) and incubation for 3 min at 16 °C. The reaction was stopped with 180 μl Stop Solution II, treated with proteinase K (0.24 mg/ml), and phenol/chloroform-extracted. After excess label was removed using a Micro Bio-Spin 6 Column (Bio-Rad), plasmids were ethanol-precipitated, and resuspended in 5 μl ELB. The labelled plasmid was used in a replication reaction and samples at various times were extracted and digested with HincII. Fragments were separated on a 0.8% denaturing agarose gel in alkaline buffer, after which the gel was dried on Amersham Hybond-XL membrane and exposed to a phosphor screen. The band intensity of the X-shape structure was measured with Typhoon TRIO+ and quantified using ImageQuant TL software. The highest value was set at 100% for the X-shape structure.

### Two-dimensional gel electrophoresis (2DGE)

2DGE was performed as described previously (Long et al., 2011). Replication samples of pICL at various times were extracted and digested with HincII. Fragments were then analyzed by 2DGE. The first-dimension gel consisted of 0.4% agarose run in 0.5x TBE buffer at 0.86 V/cm for 24 h at room temperature. The lanes of interest were cut out, casted across the top of the second-dimension gel consisting of 1% agarose with 0.3 μg/ml ethidium bromide, and run in 0.5x TBE containing 0.3 μg/ml ethidium bromide with buffer circulation at 3.5 V/cm for 14.5 h at room temperature. The gel was dried on Amersham Hybond-XL membrane and exposed to a phosphor screen. DNA was visualized using a Typhoon TRIO+.

### Plasmid pull-down assay

Plasmid pull-down assay was performed as described previously with modifications (Budzowska et al., 2015). At the indicated times, 5 μl of the pICL replication samples were mixed with 3.75 μl LacI-coupled magnetic beads (Dynabeads M-280; Life Technologies) suspended in 25 μL ELB containing 0.25 mg/ml BSA, 2.5 mM CaCl_2_, and 0.03% Tween 20, and incubated for 30 min on ice. The beads were washed 3 times with 37.5 μl ELB containing 0.25 mg/ml BSA, 2.5 mM CaCl_2_, and 0.02% Tween 20, dried, and suspended in 20 μl 1x SDS sample buffer. Plasmid-bound proteins were then separated by SDS-PAGE and visualized by western blotting using the indicated antibodies.

### Chromatin immunoprecipitation (ChIP)

ChIP was performed as described previously (Pacek et al., 2006). At the indicated times, pICL replication samples were crosslinked with ELB containing 1% formaldehyde for 10 min at room temperature. A non-related undamaged control plasmid (pQuant) was added to the replication reactions to assess background binding of the proteins. After quenching the formaldehyde by addition of glycine (125 mM final concentration), the samples were passed through a Micro Bio-Spin 6 Chromatography column (Bio-Rad), sonicated, and immunoprecipitated with 5 μg of the indicated antibodies bound to PAS beads. The protein-bound DNA fragments were eluted with ChIP elution buffer (50 mM Tris pH7.5, 10 mM EDTA, 1% SDS) and the crosslinks were reversed by incubation at 42 °C for 6 h and subsequently at 70 °C for 9 h. DNA was then phenol/chloroform-extracted for analysis by quantitative real-time PCR with the following primers: pICL (5’-AGCCAGATTTTTCCTCCTCTC-3’ and 5’-CATGCATTGGTTCTGCACTT-3’) and pQuant (5’-TACAAATGTACGGCCAGCAA-3’ and 5’-GAGTATGAGGGAAGCGGTGA-3’). The values from pQuant primers were subtracted from the values for pICL primers. Antibodies used for ChIP were purified with PAS beads.

### Quantification and statistical analysis

Statistical analysis was performed using Graphpad Prism software. Statistical significance was determined using the Kruskal-Wallis test with Dunn’s post-test.

### Data and software availability

Data presented in the paper are available from the corresponding author upon reasonable request.

---

**Figure S1 (related to Figure 1).**
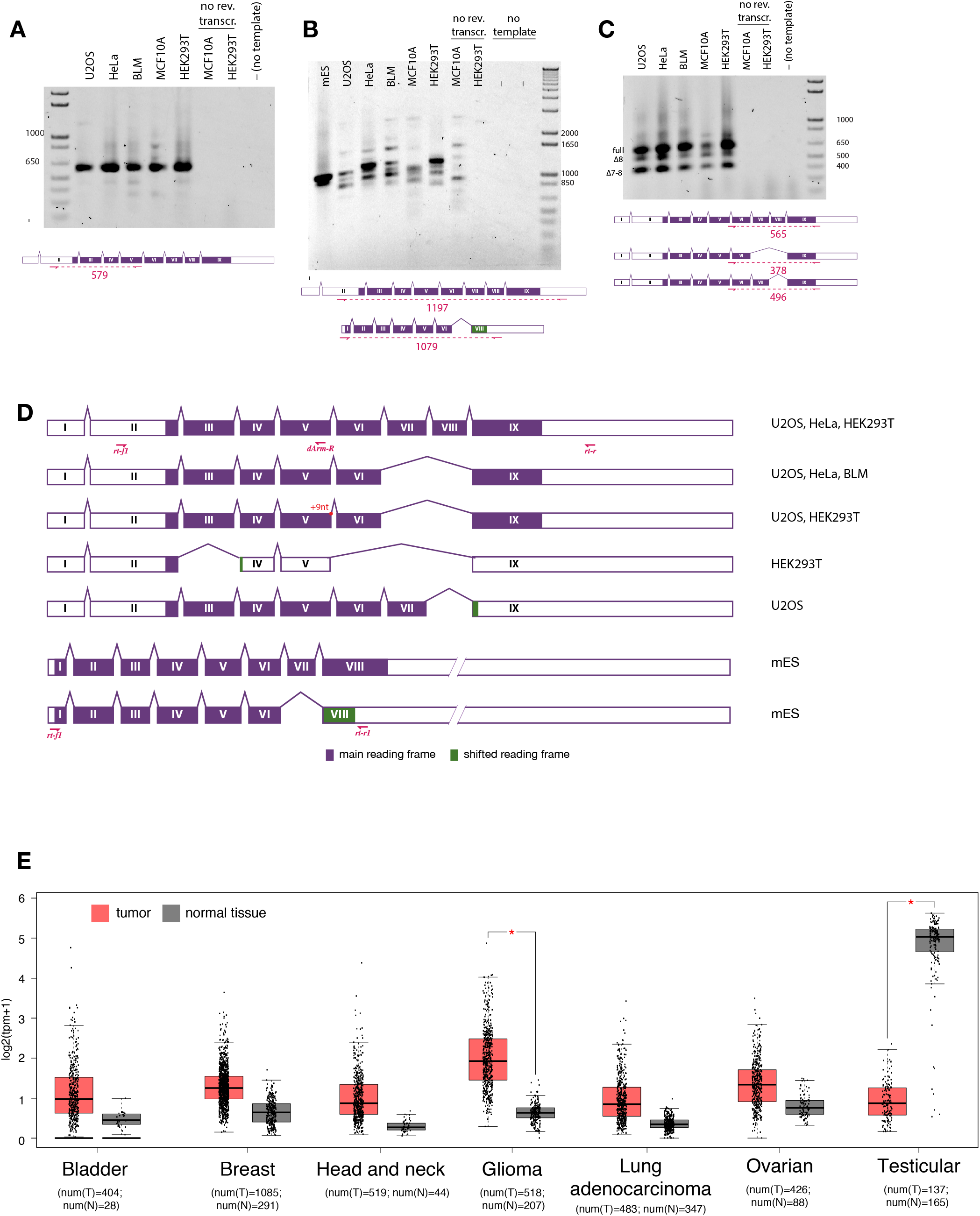
(**A-C**) PCR amplification of HSF2BP CDS fragments from first-strand cDNA produced from total RNA extracted from indicated human cancer cell lines. Primers for 5’ half (A), complete CDS (B) or 3’ half (C) were used, locations are indicated on schematic exon splicing diagrams shown below the gel images. Splice forms as identified by Sanger sequencing are depicted underneath the gel image in (C) and corresponding bands are indicated. (**D**) Schematic depiction of all HSF2BP splicing patterns deduced from the sequences of cloned PCR products amplified with the primers designed for HSF2BP CDS amplification (B) in different human and mES cell lines. Extent and splicing patterns of the regions outside of the amplicon and containing UTRs (empty boxes) were not determined and were deduced from genomic annotation. Changes in the reading frame in the coding sequence (filled boxes) is indicated by different fill color. Alternative splicing donor site leading to a 9-bp in-frame addition at the end of V in one of the splice variants is indicated. Primer annealing sites are indicated with red arrows. (**E**) Expression of HSF2BP in tumor material and normal tissues of different origin. The plot was generated from public RNA-seq data using GEPIA server (Tang et al., 2017). Number of normal (num(N)) and tumor (num(T)) samples is indicated; tpm – transcripts per kilobase million.

**Figure S2 (related to figure 2).**
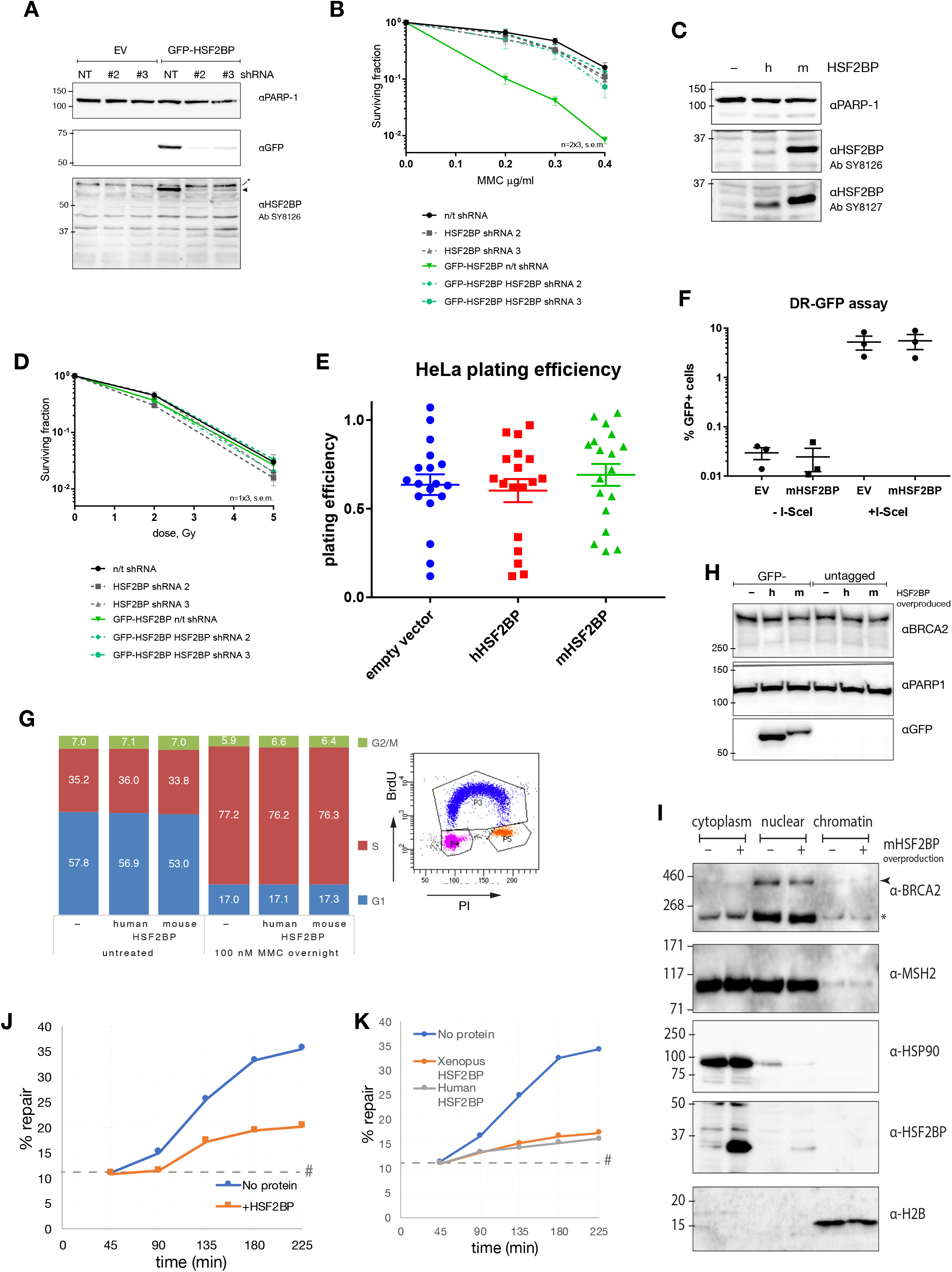
**A**) Immunoblot on the total protein extracts from cells used in the clonogenic DNA damage sensitivity assays shown in panels B and D probed with the indicated antibodies. (**B**) Clonogenic survival of U2OS cells stably expressing GFP-hHSF2BP or GFP, and either anti-hHSF2BP shRNA (#2 or #3), or non-targeting shRNA. GFP-HSF2PB U2OS clone 5 was used in these early experiments, which was produced by random integration of the CMV→GFP-HSF2BP expression construct pAZ019. Cells were treated with the indicated concentrations of MMC for 1.5h. (**C**) Immunoblots on the total protein extracts from cells used in the survivals **[Fig 2A-D]** probed with two different anti-HSF2BP sera (SY8126 and SY8127) and anti-PARP1 as loading control. (**D**) Clonogenic survival as in panel B, but instead of MMC cells were exposed to the indicated doses of γ-irradiation. (**E**) Plating efficiency of the cells from clonogenic survivals in **[Fig 2A-D]**. (**F**) Efficiency of DSB-induced HR-mediated gene conversion in U2OS cells containing direct-repeat GFP I-SceI nuclease inducible gene conversion reporter construct (Puget et al., 2005) and stably transformed with mHSF2BP overproduction or control empty PiggyBac vector (EV). Cells were lipofected with I-SceI expression construct, empty vector, or transfection efficiency reporter, and analyzed by FACS two days later. Plotted percentages of GPF+ cells that arise after HR-mediated repair of the I-SceI DSB were adjusted for transfection efficiency. (**G**) Cell cycle stage distribution in HSF2BP-overproducing or control cells with and without MMC treatment (100 nM overnight) was analyzed by FACS. For S-phase detection cells were pulsed with BrdU for 15 min, and BrdU was detected by FITC-conjugated anti-BrdU antibody, while DNA content was measured by PI staining. Gates used to quantify percentage of cells in G1, S and G2/M phases are shown on the example FACS plot on the right. (**H**) Immunoblot analysis of BRCA2 levels in control and HSF2BP (human and mouse, GFP-tagged and untagged) overproducing HeLa cells. Anti-PARP1 blotting was used to confirm equal loading. (**I**) Immunoblot analysis of BRCA2 distribution in different cellular compartments in HSF2BP-overexpressing and control HeLa cells. The efficiency of biochemical fractionation method used (Rodrigue et al., 2006) was estimated by probing for the localization of HSP90 (cytoplasm), MSH2 (nucleoplasm) and histone H2B (chromatin). Asterisk indicates non-specific band. (**J**) Repeat of the Xenopus egg extract pICL repair assay shown in **[Fig 2F]**. #, SapI fragments from contaminating uncrosslinked plasmid present in varying degrees in different pICL preparations. (**K**) ICL repair efficiency in Xenopus egg extract in the presence or absence of purified recombinant His-tagged human and Xenopus HSF2BP. #, SapI fragments from contaminating uncrosslinked plasmid present in varying degrees in different pICL preparations.

**Figure S3 (related to figure 3).**
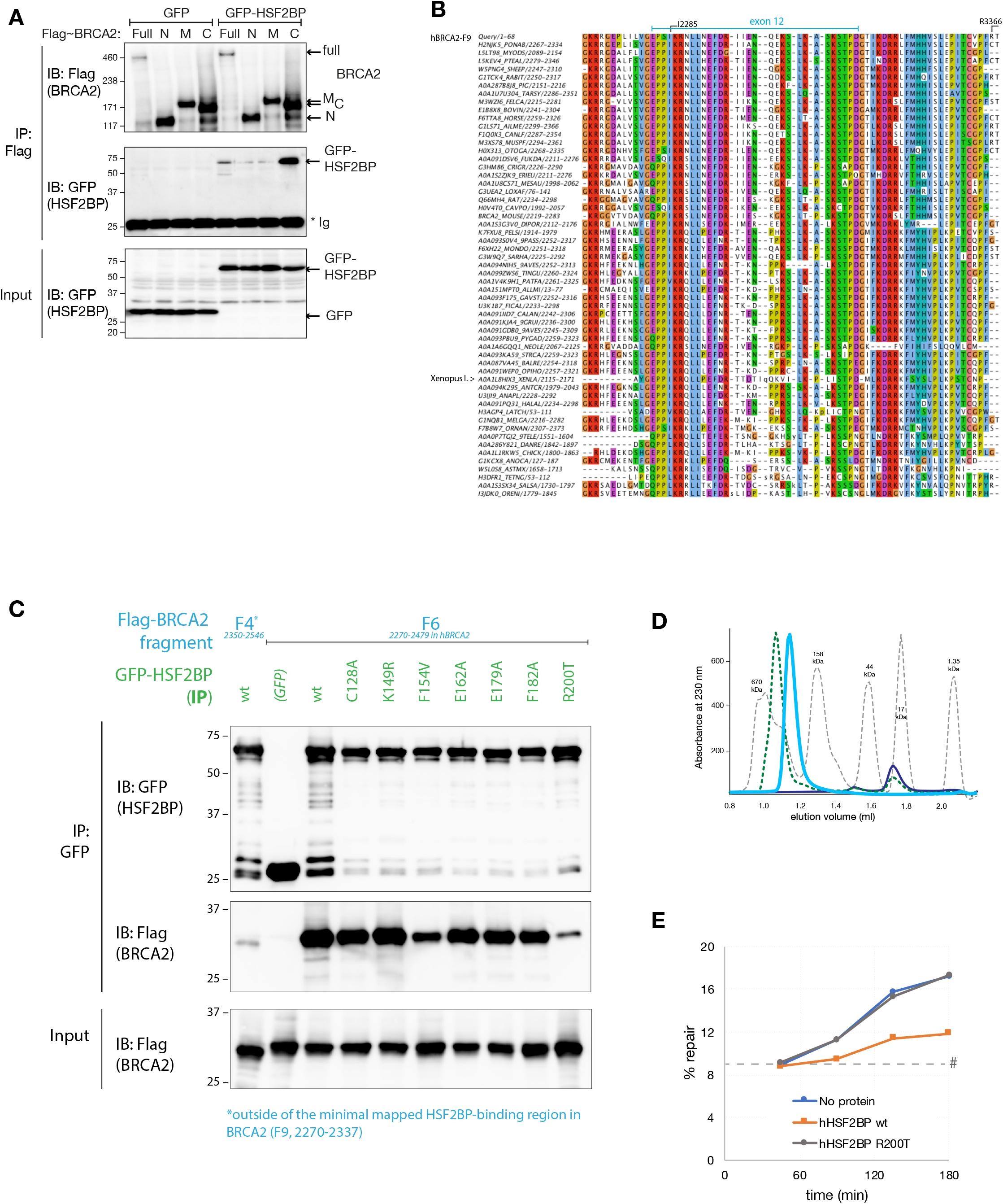
(**A**) Co-immunoprecipitation of GFP-HSF2BP with N-terminal, middle, and C-terminal fragments of BRCA2. HeLa cells stably producing Flag-BRCA2 (full- length, or the fragments) were electroporated with GFP- or GFP-HSF2BP expression constructs. (**B**) Multiple alignment of HSF2BP-binding domain (BRCA2-F9) from BRCA2 sequences from vertebrates (fish to mammals). Human HSF2BP was used a seed query for hmmsearch software. Matched sequence fragments were aligned, highly similar sequences were replaced by single representative (95% redundancy filter). Human and Xenopus laevis BRCA2 fragments are indicated. Sequence identifiers contain Uniport accession numbers and the residue range for the aligned fragment within BRCA2 sequence. (**C**) Co-immunoprecipitation with anti-GFP beads of Flag-BRCA2-F6 with variants of GFP-HSF2BP containing point mutations in the region required for BRCA2 interaction. HEK293T cells were transiently transfected with two expression constructs. (**D**) Size exclusion chromatograms shown in **[Fig 3E]** overlaid with protein size markers analyzed under the same conditions. (**E**) Repeat of the Xenopus egg extract ICL repair experiment shown in **[Fig 3F]**. #, SapI fragments from contaminating uncrosslinked plasmid present in varying degrees in different pICL preparations.

**Figure S4 (related to figure 4).**
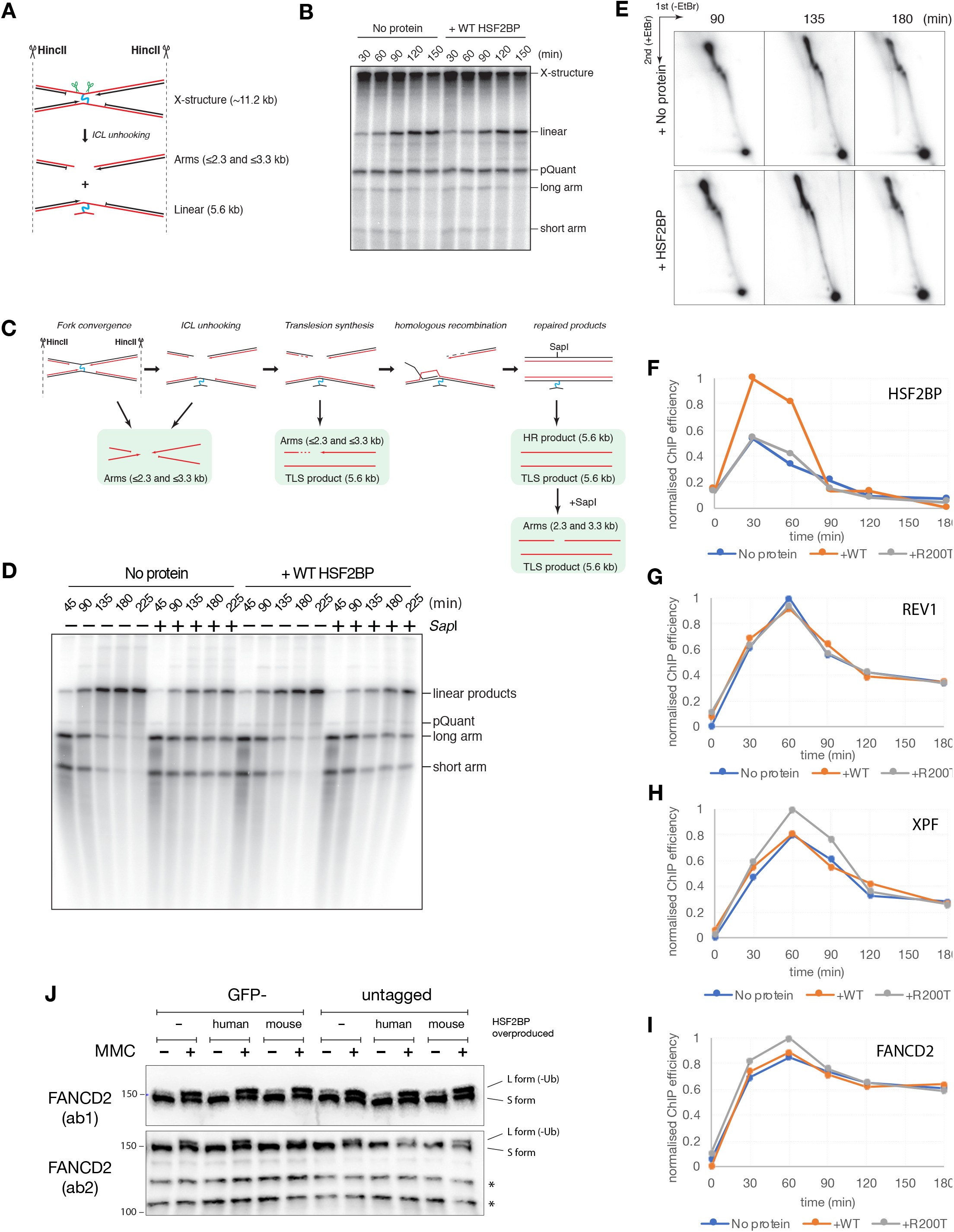
(**A**) Scheme of the lesion ICL unhooking assay. (**B**) ICL unhooking assay gel quantified in **[Fig 4A]**. (**C**) Scheme of the key events during ICL repair in Xenopus egg extract and the products measured in lesion bypass assay (see **[Fig S4D]** and **[Fig 4B]** for an example of a gel and quantification. (**D**) Lesion bypass assay gel quantified in **[Fig 4B]**. (**E**) Repeat of the experiment shown in **[Fig 4C]**. (**E-H**) Efficiency of ChIP with HSF2BP, XPF, REV1 and FANCD2 antibodies from the experiment shown in **[Fig 4F]**. (**I**) Efficiency of MMC-induced (50 ng/ml overnight) FANCD2 monoubiquitination (appearance of the slower migrating L-form) in control and HSF2BP-overproducing (human or mouse, GFP-tagged or untagged) HeLa cells. Immunoblots with two different anti-FANCD2 antibodies are shown. Asterisks indicate non-specific bands.

## References

Ameziane, N., May, P., Haitjema, A., van de Vrugt, H.J., van Rossum-Fikkert, S.E., Ristic, D., Williams, G.J., Balk, J., Rockx, D., Li, H., et al. (2015). A novel Fanconi anaemia subtype associated with a dominant-negative mutation in RAD51. Nat Commun 6, 8829.

Auerbach, A.D. (2009). Fanconi anemia and its diagnosis. Mutat Res 668, 4–10.

Béroud, C., Letovsky, S.I., Braastad, C.D., Caputo, S.M., Beaudoux, O., Bignon, Y.-J., Bressac-de Paillerets, B., Bronner, M., Buell, C.M., Collod-Béroud, G., et al. (2016). BRCA Share: A Collection of Clinical BRCA Gene Variants. Hum Mutat 37, 1318–1328.

Bièche, I., and Lidereau, R. (1999). Increased level of exon 12 alternatively spliced BRCA2 transcripts in tumor breast tissue compared with normal tissue. Cancer Research 59, 2546–2550.

Biswas, K., Das, R., Alter, B.P., Kuznetsov, S.G., Stauffer, S., North, S.L., Burkett, S., Brody, L.C., Meyer, S., Byrd, R.A., et al. (2011). A comprehensive functional characterization of BRCA2 variants associated with Fanconi anemia using mouse ES cell-based assay. Blood.

Budzowska, M., Graham, T.G.W., Sobeck, A., Waga, S., and Walter, J.C. (2015). Regulation of the Rev1-pol ζ complex during bypass of a DNA interstrand cross-link. Embo J 34, 1971–1985.

Chalkley, G.E., and Verrijzer, C.P. (2004). Immuno-depletion and purification strategies to study chromatin-remodeling factors in vitro. Meth Enzymol 377, 421–442.

Claes, K., Poppe, B., Coene, I., Paepe, A.D., and Messiaen, L. (2004). BRCA1 and BRCA2 germline mutation spectrum and frequencies in Belgian breast/ovarian cancer families. Br J Cancer 90, 1244–1251.

Cox, J., Matic, I., Hilger, M., Nagaraj, N., Selbach, M., Olsen, J.V., and Mann, M. (2009). A practical guide to the MaxQuant computational platform for SILAC-based quantitative proteomics. 4, 698–705.

Enoiu, M., Ho, T.V., Long, D.T., Walter, J.C., and Schärer, O.D. (2012). Construction of plasmids containing site-specific DNA interstrand cross-links for biochemical and cell biological studies. Methods Mol Biol 920, 203–219.

Epstein, D.J. (2009). Cis-regulatory mutations in human disease. Brief Funct Genomic Proteomic 8, 310–316.

Feng, W., and Jasin, M. (2018). Homologous Recombination and Replication Fork Protection: BRCA2 and More! Cold Spring Harbor Symposia on Quantitative Biology 035006.

Fokkema, I.F.A.C., Taschner, P.E.M., Schaafsma, G.C.P., Celli, J., Laros, J.F.J., and Dunnen, den, J.T. (2011). LOVD v.2.0: the next generation in gene variant databases. Hum Mutat 32, 557–563.

Folias, A., Matkovic, M., Bruun, D., Reid, S., Hejna, J., Grompe, M., D’Andrea, A.D., and Moses, R. (2002). BRCA1 interacts directly with the Fanconi anemia protein FANCA. Hum Mol Genet 11, 2591–2597.

Garcia-Higuera, I., Taniguchi, T., Ganesan, S., Meyn, M.S., Timmers, C., Hejna, J., Grompe, M., and D’Andrea, A.D. (2001). Interaction of the Fanconi anemia proteins and BRCA1 in a common pathway. Mol Cell 7, 249–262.

Gibson, D.G., Young, L., Chuang, R.-Y., Venter, J.C., Hutchison, C.A., and Smith, H.O. (2009). Enzymatic assembly of DNA molecules up to several hundred kilobases. Nat Meth 6, 343–345.

Harding, S.D., Armit, C., Armstrong, J., Brennan, J., Cheng, Y., Haggarty, B., Houghton, D., Lloyd-MacGilp, S., Pi, X., Roochun, Y., et al. (2011). The GUDMAP database--an online resource for genitourinary research. Development 138, 2845–2853.

Hartford, S.A., Chittela, R., Ding, X., Vyas, A., Martin, B., Burkett, S., Haines, D.C., Southon, E., Tessarollo, L., and Sharan, S.K. (2016). Interaction with PALB2 Is Essential for Maintenance of Genomic Integrity by BRCA2. PLoS Genet 12, e1006236.

Hussain, S., Wilson, J.B., Medhurst, A.L., Hejna, J., Witt, E., Ananth, S., Davies, A., Masson, J.- Y., Moses, R., West, S.C., et al. (2004). Direct interaction of FANCD2 with BRCA2 in DNA damage response pathways. Hum Mol Genet 13, 1241–1248.

Hussain, S., Witt, E., Huber, P.A.J., Medhurst, A.L., Ashworth, A., and Mathew, C.G. (2003). Direct interaction of the Fanconi anaemia protein FANCG with BRCA2/FANCD1. Hum Mol Genet 12, 2503–2510.

Inano, S., Sato, K., Katsuki, Y., Kobayashi, W., Tanaka, H., Nakajima, K., Nakada, S., Miyoshi, H., Knies, K., Takaori-Kondo, A., et al. (2017). RFWD3-Mediated Ubiquitination Promotes Timely Removal of Both RPA and RAD51 from DNA Damage Sites to Facilitate Homologous Recombination. Mol Cell 66, 622–634.e628.

Jaenisch, R., and Bird, A. (2003). Epigenetic regulation of gene expression: how the genome integrates intrinsic and environmental signals. Nat Genet 33 Suppl, 245–254.

Jensen, R.B., Carreira, A., and Kowalczykowski, S.C. (2010). Purified human BRCA2 stimulates RAD51-mediated recombination. Nature.

Kim, Y., Spitz, G.S., Veturi, U., Lach, F.P., Auerbach, A.D., and Smogorzewska, A. (2013). Regulation of multiple DNA repair pathways by the Fanconi anemia protein SLX4. Blood 121, 54–63.

Klein Douwel, D., Boonen, R.A.C.M., Long, D.T., Szypowska, A.A., Räschle, M., Walter, J.C., and Knipscheer, P. (2014). XPF-ERCC1 acts in Unhooking DNA interstrand crosslinks in cooperation with FANCD2 and FANCP/SLX4. Mol Cell 54, 460–471.

Klein Douwel, D., Hoogenboom, W.S., Boonen, R.A., and Knipscheer, P. (2017). Recruitment and positioning determine the specific role of the XPF-ERCC1 endonuclease in interstrand crosslink repair. Embo J 36, 2034–2046.

Knies, K., Inano, S., Ramirez, M.J., Ishiai, M., Surrallés, J., Takata, M., and Schindler, D. (2017). Biallelic mutations in the ubiquitin ligase RFWD3 cause Fanconi anemia. J Clin Invest 127, 3013–3027.

Knipscheer, P., Räschle, M., Schärer, O.D., and Walter, J.C. (2012). Replication-coupled DNA interstrand cross-link repair in Xenopus egg extracts. Methods Mol Biol 920, 221–243.

Knipscheer, P., Räschle, M., Smogorzewska, A., Enoiu, M., Ho, T.V., Schärer, O.D., Elledge, S.J., and Walter, J.C. (2009). The Fanconi anemia pathway promotes replication-dependent DNA interstrand cross-link repair. Science 326, 1698–1701.

Kottemann, M.C., and Smogorzewska, A. (2013). Fanconi anaemia and the repair of Watson and Crick DNA crosslinks. Nature 493, 356–363.

Krawczyk, P.M., Eppink, B., Essers, J., Stap, J., Rodermond, H., Odijk, H., Zelensky, A.N., van Bree, C., Stalpers, L.J., Buist, M.R., et al. (2011). Mild hyperthermia inhibits homologous recombination, induces BRCA2 degradation, and sensitizes cancer cells to poly (ADP-ribose) polymerase-1 inhibition. 108, 9851–9856.

Lee, T.I., and Young, R.A. (2013). Transcriptional regulation and its misregulation in disease. Cell 152, 1237–1251.

Li, L., Biswas, K., Habib, L.A., Kuznetsov, S.G., Hamel, N., Kirchhoff, T., Wong, N., Armel, S., Chong, G., Narod, S.A., et al. (2009). Functional redundancy of exon 12 of BRCA2 revealed by a comprehensive analysis of the c.6853A>G (p.I2285V) variant. Hum Mutat 30, 1543–1550.

Lombardi, A.J., Hoskins, E.E., Foglesong, G.D., Wikenheiser-Brokamp, K.A., Wiesmüller, L., Hanenberg, H., Andreassen, P.R., Jacobs, A.J., Olson, S.B., Keeble, W.W., et al. (2015). Acquisition of Relative Interstrand Crosslinker Resistance and PARP Inhibitor Sensitivity in Fanconi Anemia Head and Neck Cancers. Clin Cancer Res 21, 1962–1972.

Lomonosov, M., Anand, S., Sangrithi, M., Davies, R., and Venkitaraman, A.R. (2003). Stabilization of stalled DNA replication forks by the BRCA2 breast cancer susceptibility protein. 17, 3017–3022.

Long, D.T., Raschle, M., Joukov, V., and Walter, J.C. (2011). Mechanism of RAD51-Dependent DNA Interstrand Cross-Link Repair. Science 333, 84–87.

Long, D.T., Joukov, V., Budzowska, M., and Walter, J.C. (2014). BRCA1 promotes unloading of the CMG helicase from a stalled DNA replication fork. Mol Cell 56, 174–185.

Martinez, J.S., Baldeyron, C., and Carreira, A. (2015). Molding BRCA2 function through its interacting partners. Cell Cycle 14, 3389–3395.

McCabe, N., Turner, N.C., Lord, C.J., Kluzek, K., Bialkowska, A., Swift, S., Giavara, S., O’Connor, M.J., Tutt, A.N., Zdzienicka, M.Z., et al. (2006). Deficiency in the repair of DNA damage by homologous recombination and sensitivity to poly(ADP-ribose) polymerase inhibition. Cancer Research 66, 8109–8115.

Mezulis, S., Yates, C.M., Wass, M.N., Sternberg, M.J.E., and Kelley, L.A. (2015). The Phyre2 web portal for protein modeling, prediction and analysis. Nat Protoc 10, 845–858.

Modesti, M., Budzowska, M., Baldeyron, C., Demmers, J.A.A., Ghirlando, R., and Kanaar, R. (2007). RAD51AP1 is a structure-specific DNA binding protein that stimulates joint molecule formation during RAD51-mediated homologous recombination. Mol Cell 28, 468–481.

Murai, J., Huang, S.-Y.N., Das, B.B., Renaud, A., Zhang, Y., Doroshow, J.H., Ji, J., Takeda, S., and Pommier, Y. (2012). Trapping of PARP1 and PARP2 by Clinical PARP Inhibitors. Cancer Research 72, 5588–5599.

Pacek, M., Tutter, A.V., Kubota, Y., Takisawa, H., and Walter, J.C. (2006). Localization of MCM2-7, Cdc45, and GINS to the site of DNA unwinding during eukaryotic DNA replication. Mol Cell 21, 581–587.

Petryszak, R., Keays, M., Tang, Y.A., Fonseca, N.A., Barrera, E., Burdett, T., Füllgrabe, A., Fuentes, A.M.-P., Jupp, S., Koskinen, S., et al. (2016). Expression Atlas update--an integrated database of gene and protein expression in humans, animals and plants. Nucleic Acids Res. 44, D746–D752.

Prakash, R., Zhang, Y., Feng, W., and Jasin, M. (2015). Homologous recombination and human health: the roles of BRCA1, BRCA2, and associated proteins. Cold Spring Harb Perspect Biol 7, a016600.

Prelich, G. (2012). Gene overexpression: uses, mechanisms, and interpretation. Genetics 190, 841–854.

Puget, N., Knowlton, M., and Scully, R. (2005). Molecular analysis of sister chromatid recombination in mammalian cells. 4, 149–161.

Ran, F.A., Hsu, P.D., Wright, J., Agarwala, V., Scott, D.A., and Zhang, F. (2013). Genome engineering using the CRISPR-Cas9 system. Nat Protoc 8, 2281–2308.

Rauh-Adelmann, C., Lau, K.M., Sabeti, N., Long, J.P., Mok, S.C., and Ho, S.M. (2000). Altered expression of BRCA1, BRCA2, and a newly identified BRCA2 exon 12 deletion variant in malignant human ovarian, prostate, and breast cancer cell lines. Mol Carcinog 28, 236–246.

Räschle, M., Knipscheer, P., Knipsheer, P., Enoiu, M., Angelov, T., Sun, J., Griffith, J.D., Ellenberger, T.E., Schärer, O.D., and Walter, J.C. (2008). Mechanism of replication-coupled DNA interstrand crosslink repair. Cell 134, 969–980.

Reuter, M., Zelensky, A.N., Smal, I., Meijering, E., van Cappellen, W.A., de Gruiter, H.M., van Belle, G.J., van Royen, M.E., Houtsmuller, A.B., Essers, J., et al. (2014). BRCA2 diffuses as oligomeric clusters with RAD51 and changes mobility after DNA damage in live cells. J Cell Biol 207, 599–613.

Rodrigue, A., Lafrance, M., Gauthier, M.-C., McDonald, D., Hendzel, M., West, S.C., Jasin, M., and Masson, J.-Y. (2006). Interplay between human DNA repair proteins at a unique double-strand break in vivo. Embo J 25, 222–231.

Roy, R., Chun, J., and Powell, S.N. (2012). BRCA1 and BRCA2: different roles in a common pathway of genome protection. Nat Rev Cancer 12, 68–78.

Sato, K., Shimomuki, M., Katsuki, Y., Takahashi, D., Kobayashi, W., Ishiai, M., Miyoshi, H., Takata, M., and Kurumizaka, H. (2016). FANCI-FANCD2 stabilizes the RAD51-DNA complex by binding RAD51 and protects the 5’-DNA end. Nucleic Acids Res. 44, 10758–10771.

Schlacher, K., Wu, H., and Jasin, M. (2012). A distinct replication fork protection pathway connects Fanconi anemia tumor suppressors to RAD51-BRCA1/2. Cancer Cell 22, 106–116.

Sharan, S.K., Morimatsu, M., Albrecht, U., Lim, D.S., Regel, E., Dinh, C., Sands, A., Eichele, G., Hasty, P., and Bradley, A. (1997). Embryonic lethality and radiation hypersensitivity mediated by Rad51 in mice lacking Brca2. Nature 386, 804–810.

Shin, G., Kang, T.-W., Yang, S., Baek, S.-J., Jeong, Y.-S., and Kim, S.-Y. (2011). GENT: gene expression database of normal and tumor tissues. Cancer Inform 10, 149–157.

Stoepker, C., Faramarz, A., Rooimans, M.A., van Mil, S.E., Balk, J.A., Velleuer, E., Ameziane, N., Riele, Te, H., and de Winter, J.P. (2015). DNA helicases FANCM and DDX11 are determinants of PARP inhibitor sensitivity. DNA Repair (Amst.) 26, 54–64.

Szabo, C., Masiello, A., Ryan, J.F., and Brody, L.C. (2000). The breast cancer information core: database design, structure, and scope. Hum Mutat 16, 123–131.

Tan, S.L.W., Chadha, S., Liu, Y., Gabasova, E., Perera, D., Ahmed, K., Constantinou, S., Renaudin, X., Lee, M., Aebersold, R., et al. (2017). A Class of Environmental and Endogenous Toxins Induces BRCA2 Haploinsufficiency and Genome Instability. Cell 169, 1105–1118.e1115.

Tan, T.L., Essers, J., Citterio, E., Swagemakers, S.M., de Wit, J., Benson, F.E., Hoeijmakers, J.H., and Kanaar, R. (1999). Mouse Rad54 affects DNA conformation and DNA-damage-induced Rad51 foci formation. Curr Biol 9, 325–328.

Tang, Z., Li, C., Kang, B., Gao, G., Li, C., and Zhang, Z. (2017). GEPIA: a web server for cancer and normal gene expression profiling and interactive analyses. Nucleic Acids Res. 45, W98–W102.

Thorslund, T., Esashi, F., and West, S.C. (2007). Interactions between human BRCA2 protein and the meiosis-specific recombinase DMC1. Embo J 26, 2915–2922.

Tutter, A.V., and Walter, J.C. (2006). Chromosomal DNA replication in a soluble cell-free system derived from Xenopus eggs. Methods Mol Biol 322, 121–137.

van der Lelij, P., Oostra, A.B., Rooimans, M.A., Joenje, H., and de Winter, J.P. (2010). Diagnostic Overlap between Fanconi Anemia and the Cohesinopathies: Roberts Syndrome and Warsaw Breakage Syndrome. Anemia 2010, 565268.

Walter, J., Sun, L., and Newport, J. (1998). Regulated chromosomal DNA replication in the absence of a nucleus. Mol Cell 1, 519–529.

Wang, A.T., Kim, T., Wagner, J.E., Conti, B.A., Lach, F.P., Huang, A.L., Molina, H., Sanborn, E.M., Zierhut, H., Cornes, B.K., et al. (2015). A Dominant Mutation in Human RAD51 Reveals Its Function in DNA Interstrand Crosslink Repair Independent of Homologous Recombination. Mol Cell 59, 478–490.

Wang, X., Andreassen, P.R., and D’Andrea, A.D. (2004). Functional interaction of monoubiquitinated FANCD2 and BRCA2/FANCD1 in chromatin. Mol Cell Biol 24, 5850–5862.

Wilson, J.B., Yamamoto, K., Marriott, A.S., Hussain, S., Sung, P., Hoatlin, M.E., Mathew, C.G., Takata, M., Thompson, L.H., Kupfer, G.M., et al. (2008). FANCG promotes formation of a newly identified protein complex containing BRCA2, FANCD2 and XRCC3. Oncogene 27, 3641–3652.

Wilson, J.B., Blom, E., Cunningham, R., Xiao, Y., Kupfer, G.M., and Jones, N.J. (2010). Several tetratricopeptide repeat (TPR) motifs of FANCG are required for assembly of the BRCA2/D1-D2-G-X3 complex, FANCD2 monoubiquitylation and phleomycin resistance. Mutat Res 689, 12–20.

Wu, Y., Liao, S., Wang, X., Wang, S., Wang, M., and Han, C. (2013). HSF2BP represses BNC1 transcriptional activity by sequestering BNC1 to the cytoplasm. FEBS Lett.

Xia, B., Sheng, Q., Nakanishi, K., Ohashi, A., Wu, J., Christ, N., Liu, X., Jasin, M., Couch, F.J., and Livingston, D.M. (2006). Control of BRCA2 cellular and clinical functions by a nuclear partner, PALB2. Mol Cell 22, 719–729.

Yoshima, T., Yura, T., and Yanagi, H. (1998). Novel testis-specific protein that interacts with heat shock factor 2. 214, 139–146.

Zelensky, A.N., Kanaar, R., and Wyman, C. (2014). Mediators of Homologous DNA Pairing. Cold Spring Harb Perspect Biol 6, a016451–a016451.

Zelensky, A.N., Sanchez, H., Ristic, D., Vidic, I., van Rossum-Fikkert, S.E., Essers, J., Wyman, C., and Kanaar, R. (2013). Caffeine suppresses homologous recombination through interference with RAD51-mediated joint molecule formation. 41, 6475–6489.

Zelensky, A.N., Schimmel, J., Kool, H., Kanaar, R., and Tijsterman, M. (2017). Inactivation of Pol θ and C-NHEJ eliminates off-target integration of exogenous DNA. Nat Commun 8, 66.

Zhi, G., Wilson, J.B., Chen, X., Krause, D.S., Xiao, Y., Jones, N.J., and Kupfer, G.M. (2009). Fanconi anemia complementation group FANCD2 protein serine 331 phosphorylation is important for fanconi anemia pathway function and BRCA2 interaction. Cancer Research 69, 8775–8783.

